# Odor source distance is predictable from a time-history of odor statistics for large scale outdoor plumes

**DOI:** 10.1101/2023.07.20.549973

**Authors:** Arunava Nag, Floris van Breugel

**Affiliations:** Computer Science Engineering Department, University of Nevada, Reno; Integrative Neuroscience Program, University of Nevada, Reno; Ecology Evolution and Conservation Biology Program, University of Nevada, Reno; Department of Mechanical Engineering, University of Nevada, Reno

**Keywords:** odor plume statistics, olfactory navigation, turbulent plumes

## Abstract

Odor plumes in turbulent environments are intermittent and sparse. Lab-scaled experiments suggest that information about the source distance may be encoded in odor signal statistics, yet it is unclear whether useful and continuous distance estimates can be made under real-world flow conditions. Here we analyze odor signals from outdoor experiments with a sensor moving across large spatial scales in desert and forest environments to show that odor signal statistics can yield useful estimates of distance. We show that achieving accurate estimates of distance requires integrating statistics from 5-10 seconds, with a high temporal encoding of the olfactory signal of at least 20 Hz. By combining distance estimates from a linear model with wind-relative motion dynamics, we achieved source distance estimates in a 60×60 m^2^ search area with median errors of 3-8 meters, a distance at which point odor sources are often within visual range for animals such as mosquitoes.

## INTRODUCTION

Odor plumes serve as a crucial sensory signal for many organisms searching for food and mates [1], as well as robotic systems geared towards odor source localization [2, 3]. In outdoor environments, these plumes are transported by turbulent air currents, which break them into complex, intermittent, and time-varying patterns [4–7]. As a result, a single encounter with an odor plume does not provide direct information about how far away an odor source is. Despite the challenges imposed by this structure, a variety of organisms, including insects [8–10], crustaceans [11], fish [12], and mammals [13], excel at following intermittent odor plumes toward their source. Their success has long served as motivation and inspiration for engineers aiming to develop similarly capable algorithms for applications such as search and rescue and environmental monitoring [14–16]. How animals and robots alike can solve the challenging task of plume tracking fundamentally depends on the information they can extract from their experience, and in particular, under natural wind conditions. Here, we address a fundamental question in the field of odor plume dynamics and navigation: can source distance information realistically be extracted from odor plume statistics for an agent moving across large spatial scales under natural wind conditions?

Historically, the prevailing dogma has been that animals primarily respond to their immediate olfactory experience in a reactive manner by surging upwind after an odor encounter and casting crosswind after losing the plume [8], and many algorithms for robotic systems have taken a similar approach [17–20]. However, subsequent studies have shown that animals do modulate plume tracking decisions based on a time history of odor encounters [21, 22]. Furthermore, animals qualitatively change their behavior when they get close to the source, including flying insects [23], which approach visual features when they get close [8, 24], and dogs [25] that modulate their search speed (see also [26] for a review). These observations suggest that animals are able to extract information about source distance from their olfactory experience, either in the form of a true distance estimate or through a simple correlative heuristic. Whether or not this is possible, and which statistical features of the olfactory experience are most informative, especially over large spatial scales with natural outdoor flow conditions, is not well understood.

To describe the statistics of intermittent odor plumes we use the established term *whiff* to indicate a collection of odor encounters above some threshold (also referred to as a *bout*), and the term *blank* for anything below that threshold [7, 27]. Prior experiments suggest that both the intensity of whiffs (i.e. concentration) and their timing (e.g. whiff frequency) are correlated with distance. Celani et. al [7] showed agreement between theory and both lab and field experiments (from [28]) that the probability distributions of whiff frequencies and concentrations vary with distance for a stationary sensor. Additional experiments on spatial scales of a meter or so using experimental and computational approaches and stationary sensors have further confirmed that both timing and intensity of odor whiffs carry information correlated with distance to the source [27, 29–31]. Across these studies researchers have used a diversity of statistics to characterize whiff structure, including: mean odor concentration, whiff frequency, average slope of whiffs, average duration of blanks and whiffs, intermittency factor, and variance of whiff frequency. Many of these statistics do modulate olfactory navigation behaviors of animals such as insects: concentration modulates the probability of turning upwind in flying flies [21], and odor encounter frequency and other statistics drive olfactory navigation decisions in walking flies [32–36]. What remains unknown, is the feasibility of distinguishing between distributions of whiff statistics from a finite amount of time in natural environments, where wind can be quite variable in both speed and direction [37]. In this paper, we tested whether the timing and concentration features of an odor signal can be used to estimate distance from the source in outdoor environments at spatial scales of 60×60 m^2^. We found that features of individual whiffs have a very weak correlation with distance from the source. However, by accumulating odor signal statistics–specifically whiff concentration and duration–over a 10-second time window from a moving sensor (thus capturing both spatial and temporal information) it is possible to estimate the distance to an odor source with surprising accuracy, especially under consistent wind conditions. However, this only works well if odor information is encoded at a relatively high temporal frequency–with a low pass filter cutoff of 20 Hz or more. Intriguingly, our findings for both the encoding frequency and the temporal integration time are consistent with established observations for model plume tracking organisms such as the fruit fly.

## RESULTS

### Outdoor measurements of odor plumes

To investigate the correlation between odor signal statistics and distance to the source, we collected data in two environments. First, we selected a flat desert playa in Northern Nevada’s Black Rock Desert (Fig. 1A). We used a propylene tracer gas as an odor source, emitted from a 0.014 m diameter copper tube 2 meters above the ground at a pressure of 10 PSI. We developed a mobile sensor stack consisting of a photo-ionization detector to record odor concentration, a GPS RTK receiver to record position, and an IMU (Inertial Measurement Unit) to measure angular velocity (Fig. 1B). We placed 7 wind sensors around the odor source to measure the ambient wind direction and speed, and traversed this 60×60 m^2^ space by foot while carrying the sensor stack for 6 hours (Fig. 1C).

**Figure 1.**
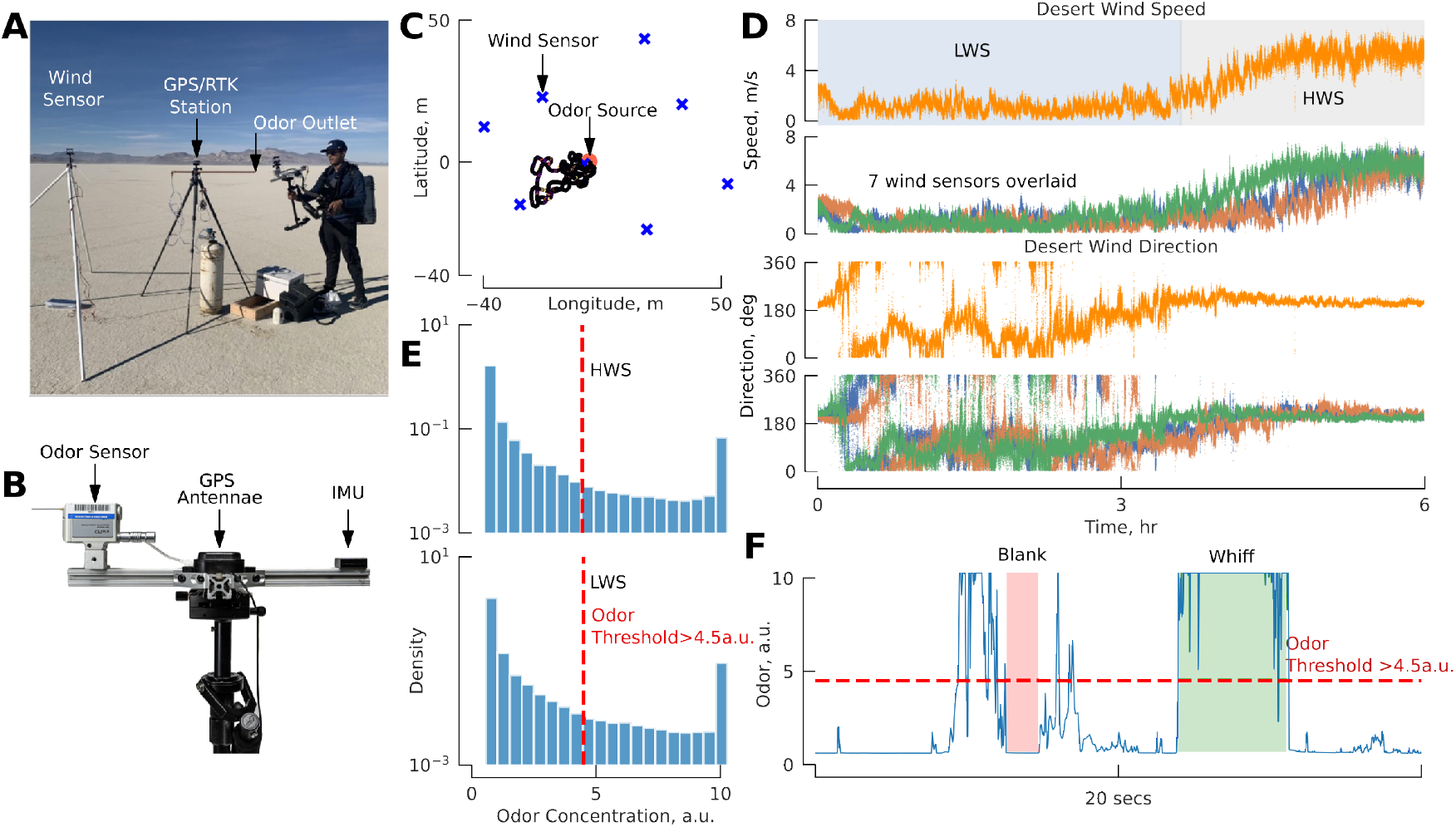
Experiment design: desert playa. **A**. Image from our Black Rock Desert field site showing the propylene odor source, GPS RTK antennae, one of seven ultrasonic anemometers, and gimbal supported sensor unit carried by one of the authors. **B**. Mobile sensor unit including a mini photo-ionization detector, GPS antenna, and IMU. **C**. Representative portion of our walking trajectory shown in relation to the odor source and anemometers. **D**. Wind speed and direction as a function of time for the duration of the 6-hour experiment for the anemometer closest to the odor source (orange traces), and all other anemometers (other colors). In our subsequent analysis we divided the data collected from this field site into two separate scenarios corresponding to the period characterized by low wind speeds and high directional variability (blue shading - HWS), and high wind speeds and low directional variability (gray shading - LWS). **E**. Log-scaled histogram of odor concentration recordings for this field site. **F**. Representative time trace of the odor signal, illustrating the intermittent nature characterized by whiffs (contiguous odor encounters greater than the 4.5 a.u. threshold) and blanks (times when the odor signal was below the threshold). More such odor traces can be seen in Fig. S3

The horizontal wind speed and direction were spatially consistent throughout the duration of the experiment (Fig. 1D), with comparatively low magnitudes of vertical wind (Fig. S1A). In subsequent analyses we used the closest stationary wind sensor to the odor source to represent the ambient wind speed and direction. Over the course of the day, the wind transitioned from low speeds and high directional variability to a strong southerly flow, indicating a shift in the local wind regime. We split our data into two datasets accordingly: “low wind speed” (LWS) < 3.5 m/s, and “high wind speed” (HWS) > 3.5 m/s (Fig. 1D).

For our second environment, we chose the Whittell Forest along Eastern Sierra Front in Nevada (Fig. 2A,B). We used a similar experimental arrangement but with the wind sensors positioned in an L-shape (Fig. 2C). Although in this environment we did find a higher degree of spatial variability of the wind, overall trends were similar across wind sensors and we again chose the closest wind sensor as a reference for the ambient wind speed and direction (Fig. 2D). Again, vertical wind speed was comparatively low (Fig. S1B).

**Figure 2.**
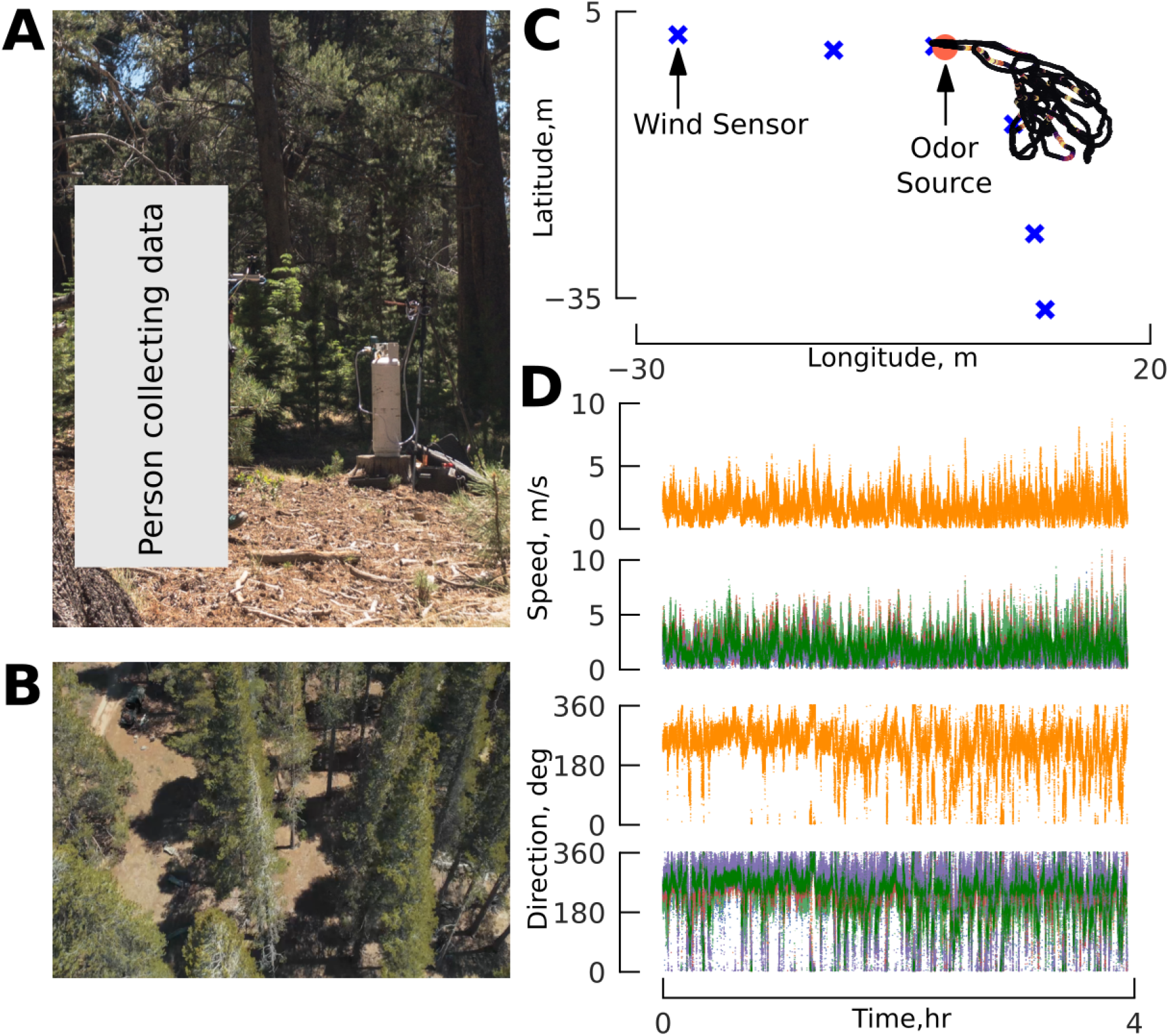
Experimental design: forest. **A**. We collected data by traversing the area surrounding the odor source on foot. **B**. Aerial view of the field site illustrating the tree density. **C**. Representative portion of our walking trajectory around the odor source, and arrangement of the ultrasonic anemometers. **D**. Wind speed and direction as a function of time for the anemometer closest to the odor source (orange traces), and all other anemometers.

The high variability in wind direction in our LWS and Forest environments would have resulted in very few encounters with the odor plume if we had used a stationary odor sensor or one guided by entirely random movements. Instead, we used our real-time odor concentration signal to drive audio cues allowing us to center our walking trajectory around the plume to efficiently gather data about as many individual odor whiffs as possible. We attempted to follow trajectories reminiscent of the casting motifs exhibited by flying insects [8]. The majority of our movement was in the plane horizontal to the ground at walking speeds of 1-2 m/s. Our changes in direction in line with the source, and perpendicular to the source, ranged from 0.02-0.2 Hz (Fig. S2). For comparison, flies and moths cast at about 0.5-1 Hz=1-2 turns/sec [8, 38, 39]).

We made small vertical movements on the scale of *±*0.5 m, generally staying aligned with the altitude of the source. Encounters with odors were infrequent: most of the time, our photo-ionization detector recorded values below 2 a.u., with occasional spikes in the readings (Fig. 1E). The miniPID sensor recorded measurements between 2 a.u. and 4 a.u. in the field when testing only ambient wind or human breath. We established a threshold of 4.5 a.u. (2.2 times the standard deviation [34] (*σ*) of all odor measurements) to categorize odor signals as either *whiffs* or *blanks* (Fig. 1F)to minimize ambient noise interference, drawing inspiration from the techniques used in the study by Jayaram et al. (2021). We also performed a sensitivity analysis to quantify the impact of different thresholds on our overall conclusions (Fig. S4).

We made our outdoor plume measurements on two days that were a part of our larger survey of near surface wind characteristics [37]. The ambient wind conditions for our odor plume experiments spanned close to the full range of typical conditions that we observed across 10 days of wind measurements in playa, sage steppe, and forest environments (Fig. S5). Altogether, our plume measurements from the desert and forest experiments spanned 10 hours of active data collection resulting in a total of 11,107 whiffs (see Table S1 for breakdown across scenarios).

### Distribution of odor whiff locations is driven by wind characteristics

Before analyzing the correlation between odor statistics and distance to the source, we wanted to visualize the shape of the odor plumes in each condition. Because the wind direction varied throughout the course of our experiments, we developed an approach to “straighten” the data by aligning encounters relative to an ideal streakline. We determined the ideal streakline by integrating the wind velocity vectors over time. Figure 3A shows six snapshots of this moving streakline evolving over time. For each moment in time, we determined the shortest distance from the sensor location to the streakline centerline, and the distance from the sensor to the source. These two values allowed us to project sensor locations into a new streakline-centered coordinate frame (Fig. 3B).

**Figure 3.**
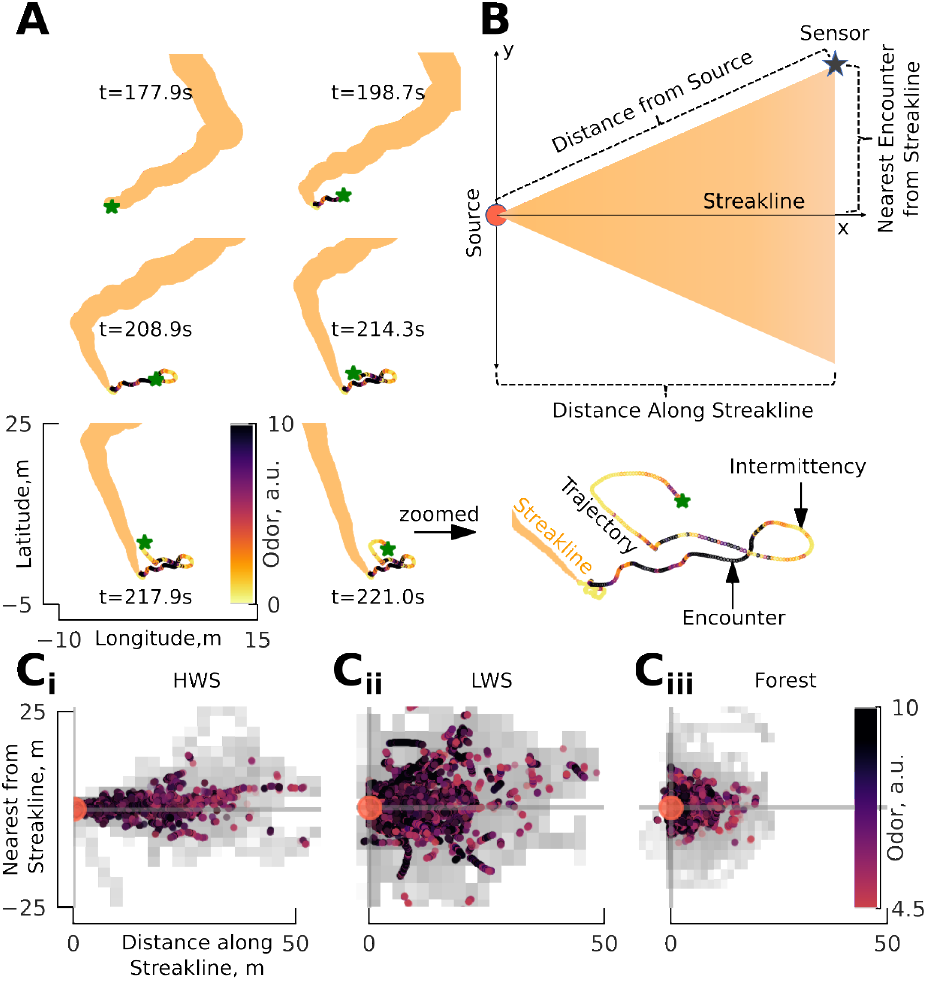
Streakline-aligned odor encounters reveal approximate shape and extent of the outdoor odor plumes. **A**. Six snapshots spanning 43 seconds showing the ideal streakline (orange, determined by integrating the ambient wind vector), together with our walking trajectory color-coded by the odor concentration recorded at each location. Green star indicates the location of the sensor for the same moment that the streakline is shown. The width of the streakline is for illustration purposes, not drawn to scale. **B**. To visualize the extent of the plumes despite the time varying wind direction we transformed each point along the trajectory from latitude/longitude to streakline coordinates. We calculated the distance from the sensor to the source and the shortest distance to the streakline. **C**. Using the approach described in **B**, we transformed each whiff (i.e. when the odor signal was greater than 4.5 a.u.) into streakline coordinates for each of the three scenarios (i-iii). Gray shading indicates time spent in streakline coordinate space, where darker shade indicates time spent more than 10 seconds. For a more clear histogram of time spent please refer to Fig. S6. N=2,842; 3,522; 4,986 whiffs for HWS, LWS, and Forest, respectively.

Figure 3C shows the streakline-centered locations where we encountered odors for all three scenarios. In the high wind speed (HWS) dataset from the desert playa, the plume was quite narrow with odor encounters occurring close to the ideal streakline, and becoming rare at distances of 25 m away (Fig. 3C_*i*_). For the low wind speed (LWS) dataset, odor encounters were spread out away from the streakline center much more widely (Fig. 3C_*ii*_). For our forest dataset (Fig. 3C_*iii*_), odor encounters were mostly found within a smaller 10-15 meter radius from the source and did not have the same cone-like shape seen in the desert datasets. This could be because the wind was more variable due to the trees spread throughout the area. In each case, examples of both low and high concentration encounters can be found across the extent of the plume, though high concentration encounters were more frequent closer to the source.

### Odor whiff statistics for estimating distance to source

To test if odor signal statistics are correlated with distance from the source, we calculated the following odor statistics (Fig. 4): whiff duration (wd), mean concentration during a whiff (wc), whiff frequency (wf, following calculations from [32], see Materials and Methods), whiff moving average (wma), and whiff standard deviation (wsd) of the encounters during a whiff. We define a single (average) value for each statistic for each individual whiff, resulting in the stepwise constant plots shown in Fig. 4B-F.

**Figure 4.**
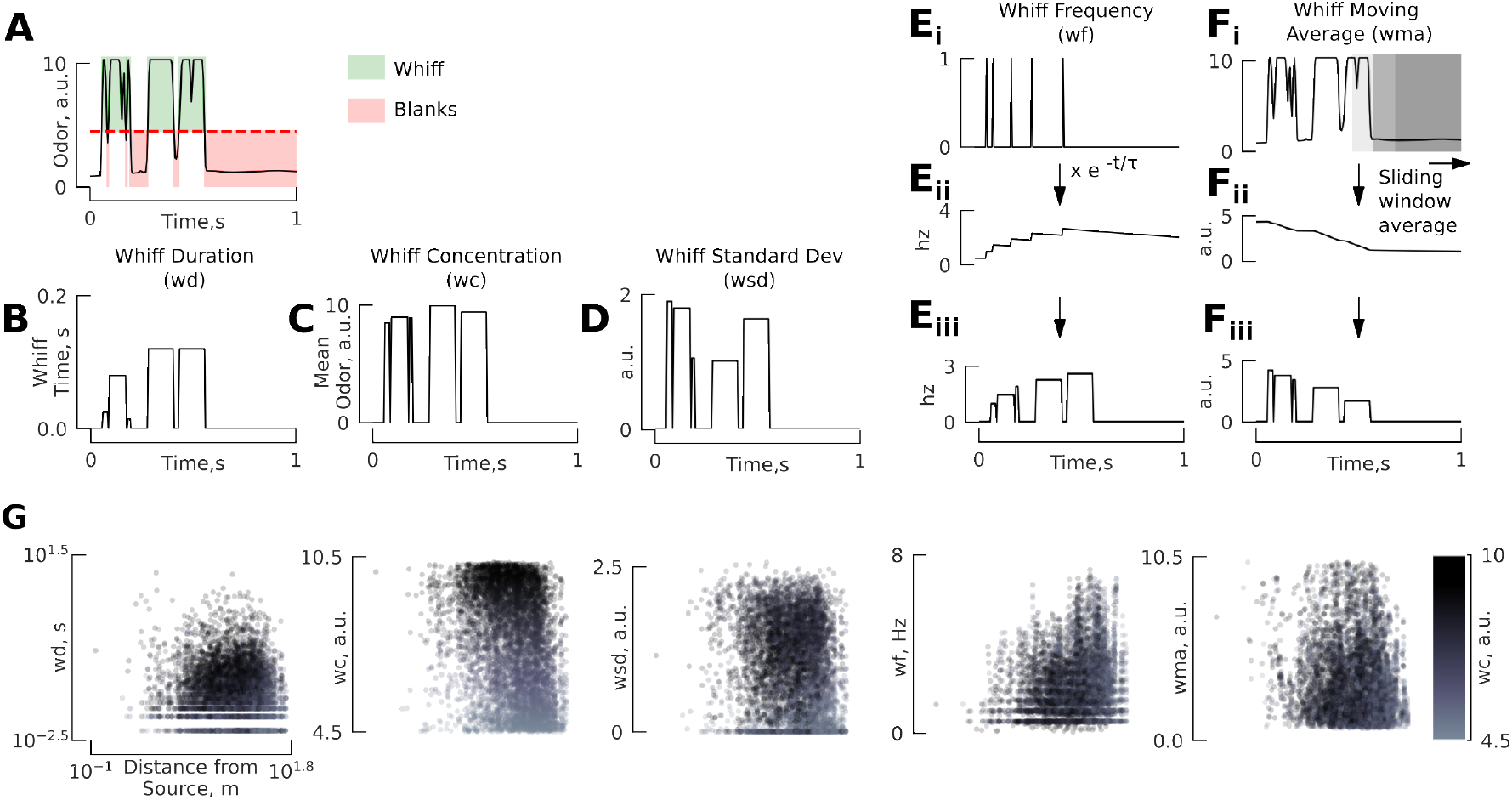
Individual whiff statistics are not strongly correlated with distance. **A**. Representative trace of odor concentration over time, color coded according to whiffs and blanks. **B-D**. For each individual whiff we calculate a single value corresponding to the whiff duration (wd), mean whiff concentration (wc), and standard deviation of the concentration within a whiff (wsd), resulting in piecewise constant time traces. **E**. Whiff frequency (wf) is calculated by convolving an exponential decay kernel (*τ* = 2) with a binary delta function corresponding to the onset of each whiff. Each whiff is assigned a single value for the whiff frequency. **F**. We calculated the moving average of odor concentration using a sliding window (1 sec), and then assigned each whiff a single value corresponding to the mean of the moving average signal during the time frame of the whiff (wma). **G**. Statistics of individual whiffs are not strongly correlated with distance to source. Each panel shows a scatter plot of the whiff statistics defined above for all three scenarios combined. Color encodes the whiff concentration. Whiff statistics for the three scenarios are shown separately in Fig. S7. N=2,842; 3,522; 4,986 whiffs for HWS, LWS, and Forest, respectively.

### Individual whiff statistics are not strongly correlated with distance

To identify potential correlations between our selection of odor statistics and distance from the source, we constructed a multiple linear regression model for each of the three scenarios (Table 1, see Fig. S8 for residual analysis). We found whiff concentration and frequency to be the most significant statistics, but both had very subtle correlations with distance. Because we collected data by moving the sensor in a biased manner, we also considered the effects of our motion relative to the instantaneous wind direction, but did not find any meaningful difference in the results (Table S2).

**Table 1.** ***R***^**2**^ and semi-partial r for Distance**∼** Odor Statistics Note: * for 1*e*^∼ 5^ < *p* < 0.05, ** for *p* < 1*e*^∼ 5^.

| Dataset | $R^2$ | semi-partial r (% contribution) | | | | |
| --- | --- | --- | --- | --- | --- | --- |
|  |  | wc | wf | wd | wma | wsd |
| HWS | 0.14 | 44.03 ** | 29.17 ** | 5.02 * | 3.41 * | 5.45 ** |
| LWS | 0.13 | 70.53 ** | 7.98 ** | 26.72 ** | 0.03 * | 0.48 * |
| Forest | 0.08 | 81.29 ** | 1.97 * | 22.10 ** | 1.21 * | 0.21 * |
| Combined | 0.13 | 89.44 ** | 10.03 * | 24.87 ** | 1.43 * | 1.37 * |

Although we found individual whiff statistics to be poorly correlated with distance, the distributions of these statistics (Fig. 4G) do exhibit consistent patterns in relation to distance. For example, whereas concentration of an individual whiff might be high, or low, across a wide range of distances, the distribution of whiff concentrations clearly trends to smaller values at higher distances. At larger distances whiff frequencies were more likely to be higher, thus high frequency but low concentration seqeuences of whiffs should be expected at large distances. At shorter distances, whiffs were rare (low whiff frequency), but when they did occur, they had a relatively long whiff duration and high whiff moving average. Finally, the overall pattern of the whiff standard deviation bears some resemblance to the whiff concentration, a relationship we explore more later. In the next section, we examine how the distribution of a time history of whiffs is correlated with distance.

### A short time history of whiff statistics reveals strong correlations with distance

In this section, instead of considering individual whiff statistics, we collect all the whiffs that occurred within a short “lookback” time window (i.e. a trajectory snippet, see Methods). This analysis results in a (sparse) probability distribution associated with each whiff statistic for a given distance (the average distance for that trajectory snippet). To determine whether these probability distributions are correlated with distance we quantified these distributions using five meta statistics:

mean (*µ*), standard deviation (*σ*), *min, max*, and kurtosis (*k*). Applying these five meta statistics to our five whiff statistics results in 25 features. We first looked for correlations between these 25 features and distance to the source for a range of lookback times. We found a sharp increase in *R*^2^ for lookback times from 1-5 seconds, with a continued but modest increase for lookback times up to 10 seconds resulting in final *R*^2^ values at 10 seconds ranging from 0.38 to 0.82 (Fig. 5A, Table 2). Figure 5B-F shows how the mean of the five whiff statistics are correlated with distance, revealing that the mean of whiff concentration and mean of the standard deviation of each whiff both exhibit significant and strongly predictive correlations (Table S3). Figure S4 shows a sensitivity analysis of these correlations to the choice of odor threshold. Across a range of thresholds, the mean of whiff concentration and the mean of whiff standard deviation consistently show the highest correlation with source distance across all three scenarios.

**Table 2.**
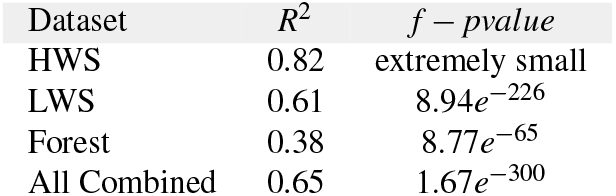
***R***^**2**^ and model significance (p-value) correlating distance from the source with all 25 meta whiff statistics collected over a lookback window = 10 seconds.

| Dataset | $R^2$ | $f - pvalue$ |
| --- | --- | --- |
| HWS | 0.82 | extremely small |
| LWS | 0.61 | $8.94e^{-226}$ |
| Forest | 0.38 | $8.77e^{-65}$ |
| All Combined | 0.65 | $1.67e^{-300}$ |

**Figure 5.**
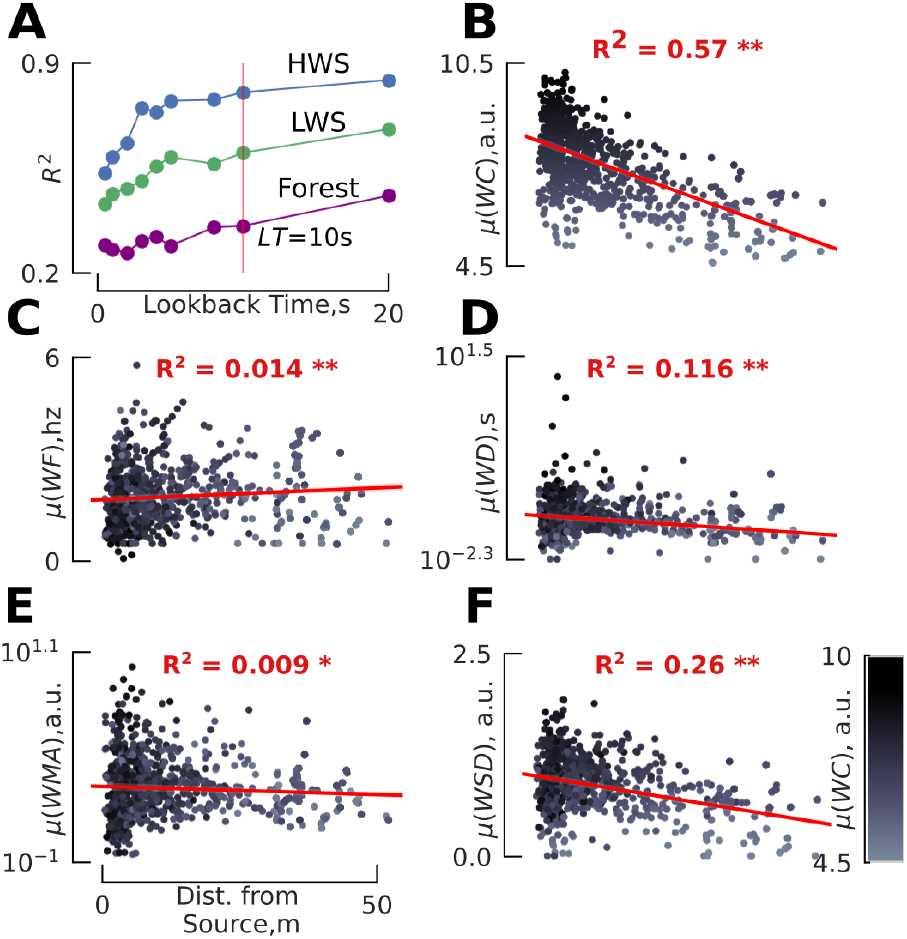
Meta statistics for whiffs occurring within a 10 second “lookback” time window are strongly correlated with distance from the source. **A**. For the collection of whiffs occurring within a chosen lookback time for randomly sampled trajectory snippets (see text) we calculated five meta statistics: mean, standard deviation, min, max, and kurtosis of each of our whiff statistics. The correlation coefficient for a linear regression relating these 25 meta statistics to the source distance rises with increasing lookback times. In subsequent analyses we chose to focus on a lookback time of 10 seconds. **B**-**F**. Distribution of the mean of each whiff statistic with respect to the distance from the source for a lookback time of 10 seconds. Here data are combined across all three scenarios. Red line shows the correlation, see Table S3 for detail.

### Concentration and timing meta statistics are correlated with source distance

Next, we set out to understand which meta statistics of our whiff statistics provide the most predictive power for distance from the source. We chose to focus on a lookback time of 10 seconds for the rest of our analysis, as beyond this point *R*^2^ does not substantially improve. We combined datasets from desert and forest environments and individually correlated the 25 meta statistics with distance (Fig. S10). To compare the relative quality of different models and avoid overfitting we used Akaike’s Information Criterion (*AIC*) [40] to construct a series of statistical models with increasing complexity (Table S4). *AIC* penalizes models with more parameters, while rewarding those that fit the data well. The single most informative meta statistic was the mean whiff concentration. Small improvements in *R*^2^ and *AIC* were realized as more meta statistics were included in the model, but after approximately 4 meta statistics the changes in *AIC* (△*AIC*) resulted in diminishing returns. With 4 parameters, the lowest *AIC* identified *µ*(*WC*), *σ* (*WD*), *max*(*WMA*), and *σ* (*WMA*) as the most informative meta statistics for estimating distance.

Notably missing from the list of meta statistics selected by our *AIC* analysis is *µ*(*WSD*), which is strongly correlated with distance (Fig. 5F). This is because *µ*(*WC*) and *µ*(*WSD*) are mutually correlated (Fig. S9, *R*^2^ = 0.51, *p* = 4.61*e*^∼ 228^), so including *µ*(*WSD*) in a statistical model that already contains *µ*(*WC*) does not provide enough novel information and would violate model assumptions, increase bias and skew the regression results. The strong correlation between *µ*(*WC*) and *µ*(*WSD*) suggests that these two meta statistics are somewhat interchangeable.

To compactly visualize all strongly correlated groups of meta statistics we constructed a hierarchically clustered matrix of the correlations between each pair of meta statistics (Fig. 6). The resulting adjacency matrix identifies *µ*(*WC*) and *µ*(*WSD*) as their own cluster (Fig. 6A), suggesting that these interchangeable statistics provide unique information compared to other whiff statistics, and that the mean and variability (*WC* and *WSD*, respectively) within a whiff are related. Both meta statistics decrease with increasing distance (Fig. 5B,F), perhaps because at larger distances diffusion has had a chance to both even out the odor distribution and reduce the concentration within a whiff. Overall this cluster can be summarized as one that captures concentration information.

**Figure 6.**
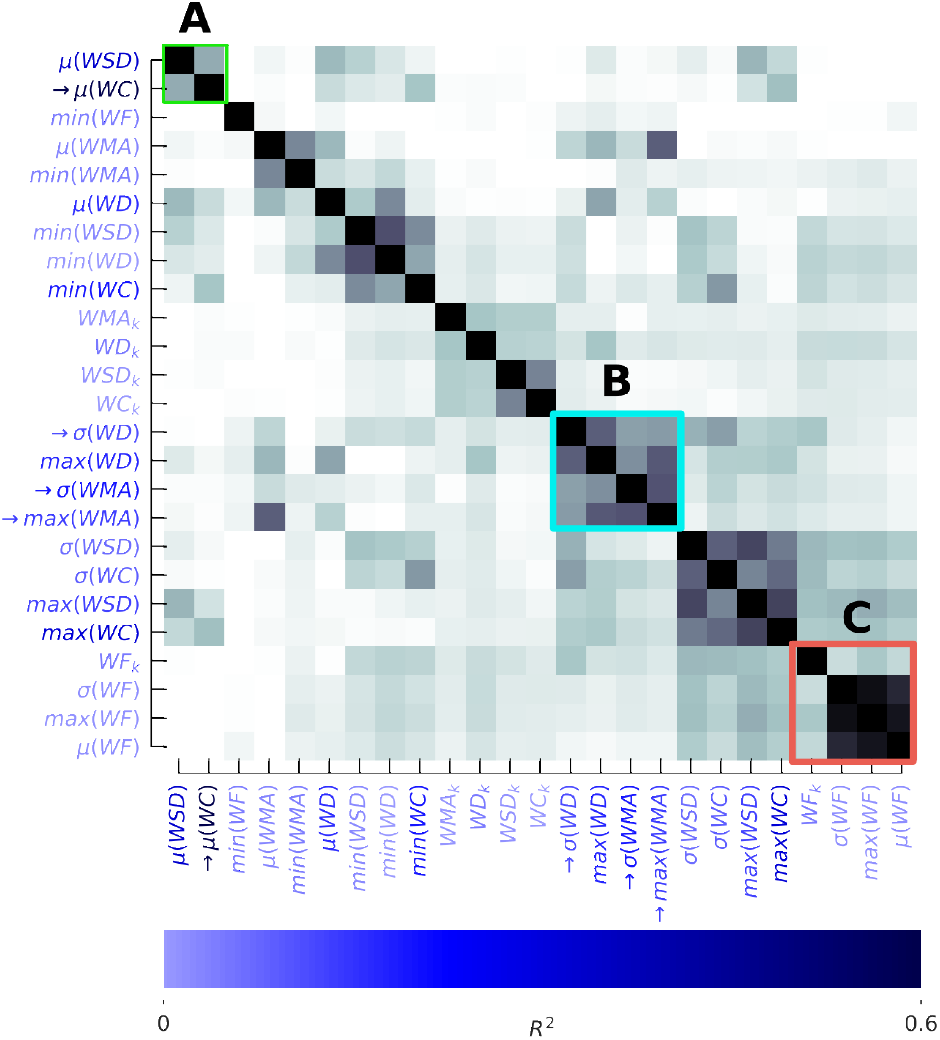
Hierarchical clustering reveals distinct clusters of mutually related meta statistics of whiffs within a lookback window. Darker colors in the adjacency matrix correspond to higher pairwise correlations between meta statistics. The *R*^2^ correlation coefficient between each individual meta statistic and distance to the source is encoded in the blue shading of the meta statistic labels. Arrows indicate meta statistics selected by the 4-parameter model from Table S4. **A** Concentration related parameters *µ*(*WC*) and *µ*(*WSD*) are mutually correlated (see also Fig. S9), and are both strongly correlated with the source distance (see also Fig. S10A). **B**. The parameters related to whiff duration – *σ* (*WD*), *max*(*WMA*), and *σ* (*WMA*) – form a distinct group, three of which are also selected by the *AIC* analysis. **C**. Whiff Frequency meta statistics form a cluster and provides unique information about the distance to the odor source, albeit low correlation with distance to the source.

Our *AIC* analysis also identified *σ* (*WD*), *max*(*WMA*), and *σ* (*WMA*) as informative, and we find that these whiff statistics form their own cluster in the adjacency matrix (Fig. 6B). Because longer whiff durations will typically result in larger whiff moving averages, this cluster can be summarized as one that captures variability in whiff duration (and its maximum), e.g. a measure of odor timing.

The next clear cluster (below and right of B) includes meta statistics that describe the variability of both whiff concentration and standard deviation across whiffs. Although these meta statistics are correlated with distance, our *AIC* analysis suggests variability of these concentration related statistics may not be as informative as their mean (note the correlation between this cluster and cluster A).

Finally, we see that meta statistics of whiff frequency form a distinct cluster (Fig. 6C), indicating that whiff frequency provides unique odor timing related information compared to our other whiff statistics. However, whiff frequency is not strongly correlated with distance in our experiment (Table S3, *R*^2^ = 0.014, *p* = 4.0*e*^∼ 06^). We discuss this apparent conflict with existing literature in the discussion.

### Integrating whiff statistics with relative motion can provide a continuous estimate of source distance

The intermittent nature of odor whiffs means that any estimate of distance based solely on whiff statistics will also be highly intermittent, and therefore of limited utility. To illustrate how this signal could be used to generate a continuous estimate of distance we used a constant velocity Kalman smoother to integrate two measurements (see Materials and Methods).

First, we used our 4-parameter statistical model for a lookback time of 10 seconds (Table S4) to generate intermittent distance predictions. We combined the data from all three of our scenarios, and for our odor delivery paradigm across these environments the distance predictions are given by the following equation,

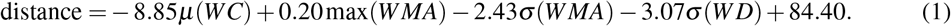

Second, we used continuous estimates of the sensor ground velocity (acquired via GPS) component parallel to the instantaneous ambient wind direction. Despite the model simplicity, our smoothed continuous distance estimates produced surprisingly accurate distance estimates in both the desert HWS and LWS scenarios, achieving median errors in absolute distance of 3.9 m and 6.26 m, respectively (Fig. 7A-B). For the Forest scenario, our estimates consistently overestimated the true distance, resulting in a median error of 7.76 m (Fig. 7C). Overall these results were relatively independent of the choice of odor threshold (Fig. S4A). For the HWS case, a threshold of 4.5 a.u. yielded the minimum error for HWS, however the errors stayed mostly constant for LWS and Forest across thresholds.

**Figure 7.**
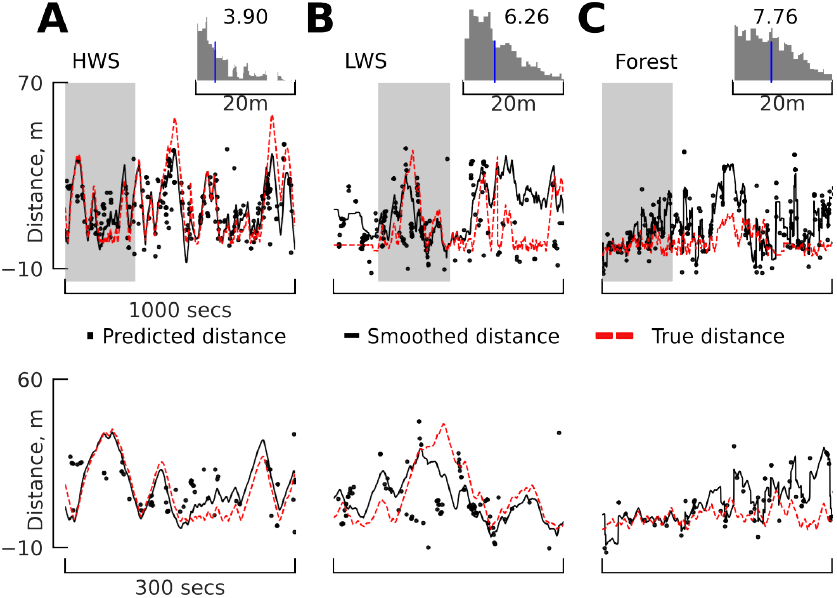
Integrating whiff statistics with relative motion in a Kalman smoother yields continuous estimates of source distance. **A-C** Upper row: Comparison of predicted, Kalman smoothed, and true distances for three scenarios. Our Kalman smoother combined intermittent distance estimates from our combined desert+forest linear model with continuous estimates of our ground velocity component parallel to the ambient wind. Histograms show the median error between the Kalman smoothed and true distance. Lower row: a magnified view of the shaded regions from the upper row. For an even zoomed-in presentation of the above smoothed results and if environment-specific prediction models are used, refer to Fig. S12.

Next we asked how much the statistical models varied across our three scenarios, and whether using environment specific models might improve the accuracy of distance estimates. Only the intercept of the model changed substantially across the scenarios, dropping consistently with increasing variability in wind direction (Fig. S11). In other words, equivalent whiff statistics in scenarios with higher wind variability generally correspond to shorter distances. Although incorporating these scenario-specific model parameters does improve the accuracy of distance estimates, the changes in performance are modest (Fig. S12).

### Odor Signal Temporal Resolution needs to be upwards of 20Hz for accurate distance estimation

Finally, we asked whether a high temporal resolution of the odor signal is necessary to accurately estimate the distance to the source, or if slower time-averaged measurements would perform equally well. To answer this question, we low-pass filtered our 200 Hz odor signal time series using a 2nd order Butterworth filter [41]. Figure 8A shows representative time traces of the filter’s effects. For each cutoff frequency, we re-ran our analysis pipeline to identify the new whiffs, their statistics, the meta statistics for a 10-sec lookback window and calculated the *R*^2^. Our results show that cutoff frequencies of less than 20 Hz generally result in substantially reduced predictive power (especially in the HWS and Forest scenarios) (Fig. 8B).

**Figure 8.**
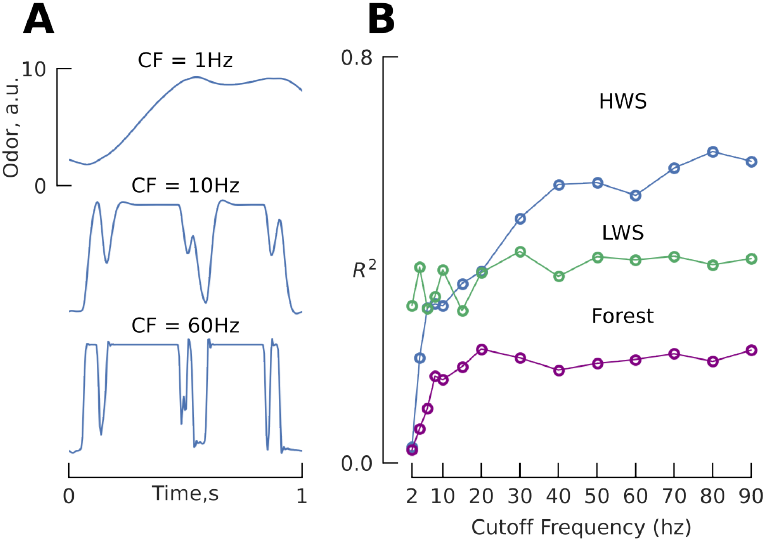
Low pass filtering odor signal degrades correlations between whiff statistics and source distance. **A**. Sample time traces of odor signal for selected cutoff frequencies used in a 2nd order Butterworth filter and varied the cutoff frequency from 1 Hz to 90 Hz (approximately half the Nyquist frequency), while keeping the order of filter constant at two. **B**. The correlation coefficient for distance correlated with *AIC* filtered whiff meta statistics for a lookback time of 10 seconds as a function of cut-off frequency (CF). Correlation coefficients shown here are small than those in Fig. 5 because this analysis uses only 4 of the 25 parameters, selected by our *AIC* analysis.

## DISCUSSION

Our outdoor experiments showed that a moving agent could estimate source distance by integrating a 5-10 second time history of whiff statistics together with their movement relative to the wind. Furthermore, we showed that this is only feasible given a high odor sampling frequency of 20 Hz or more. Consistent with prior work using meter-scale fluid dynamics simulations [27], we found that both concentration features (mean concentration and standard deviation of concentration within a whiff, Fig. 6A), and timing features (whiff duration and moving average, Fig. 6B) provide complimentary information. Contrary to wind tunnel experiments [7, 31], however, we found that whiff frequency formed its own distinct cluster which was not strongly correlated with distance (6C). Only for individual whiffs in consistent wind (our HWS case) was whiff frequency more predictive compared to whiff duration, though its utility was still limited (*R*^2^ = 0.04). In our lookback analysis whiff frequency did drop to very low values close to the source (possibly driven by either sensor movements or changes in wind direction), suggesting that perhaps a low frequency of encounters could be indicative of approaching the source. Together, our results are consistent with the idea that for a sensor moving slowly relative to wind, whiff frequency may carry useful information. However, under more naturalistic conditions that involve either dynamic wind, or a moving sensor that may or may not stay within the odor plume envelope, whiff frequency may be an unreliable cue when accumulating evidence over a longer lookback window, and the duration of individual whiffs may be more informative.

Our field experiments were necessarily limited due to the logistics of gathering this type of data. Accordingly, we chose to use a single source geometry, odorant, release concentration, and flow rate for all three scenarios. The specific values of the coefficients relating distance to each of the whiff statistics we considered would likely change for different choices of these parameters. We used a PID portable chemical sensor and utilized the system’s maximum settings, however, we did not account for scenarios of signal saturation, which could be mitigated in the future by using a sensor with a higher range, or collecting data at a variety of gain settings. Our sensitivity analysis (Fig. S4), however, suggests that the intermediate threshold we chose for our analyses (4.5 a.u.) was a near optimal choice for this dataset.

Thresholds below 2-4 a.u. led to poorer correlations with source distance (Fig. S4B-D), likely because of higher levels of ambient noise, whereas higher thresholds also resulted in lower correlations in many cases, likely by removing too much signal information. The minor improvement in distance estimation accuracy we achieved when considering environment specific models instead of a single combined model, however, suggests that distance estimation can be relatively robust to specific characteristics of the wind environment (Fig. S12). The coefficient related to the whiff concentration is likely the one that would change the most for other types of source conditions. Our hierarchical clustering analysis suggests that robustness to differences in release concentrations may be realized by relying on the standard deviation within a whiff more so than the concentration. Alternatively, a priori knowledge of the type of source might be needed to achieve reliable distance estimates. In the context of olfactory navigation among animals or machines this could be achieved through niche- or task-specific models. For example, a male moth following a sparse pheromone plume could utilize a different distance estimation model compared to a mosquito following a CO_2_ plume to a human host, or a fruit fly searching for decaying fruit matter.

Might animals, such as flying insects, use whiff statistics to estimate the distance to a source? The qualitative changes exhibited by flying insects when they get close to a source [23, 26] indicate that there is likely at least a correlational mechanism linking their olfactory experience with their behavioral decision to approach visual features [8, 24]. In our experiments we achieved an accuracy of 3-8 meters, depending on the environment. This is in line with the range over which insects such as mosquitoes are likely to begin visually resolving potential odor sources [42, 43], making it a practically useful estimate for gating such behavioral decisions.

Gating a qualitative behavioral change could be accomplished by an internal estimate of distance, or a simpler binary threshold heuristic. In either case, our results suggest that a working memory of∼ 5-10 seconds and a sampling frequency of 20 Hz are required. Experiments with flying mosquitoes [42], and flying [8] and walking fruit flies [36] indicate that odor can influence their behavior on time scales of 10 seconds. Furthermore, when flies encounter a plume multiple times they adjust the strength of their upwind turns, suggesting that they do integrate odor information across multiple encounters on timescales of at least 3 seconds [21]. Meanwhile, peripheral olfactory receptors in fruit flies are characterized by a low pass filter frequency of roughly 10-20 Hz [44, 45], and arguably even faster [46]. Finally, many of the whiff statistics we identified are known to modulate aspects of plume tracking behavior in flies: concentration regulates the strength of flying flies’ upwind turns [21]; duration has a subtle effect on flying flies’ probability of turning upwind and approaching visual features [8]; walking flies adjust their decisions based on whiff frequency [32]; and flying moths require intermittent odor signals to maintain an upwind surging flight behavior [10].

Our field experiments provide clear evidence that meaningful source distance information can be extracted from a time history of odor encounter statistics in outdoor scenarios with natural wind. Future studies with plume tracking animals should focus on answering the question of whether they use the types of statistics we identified to either generate a continuous representation of distance, or drive a binary heuristic. Meanwhile, engineers aiming to develop odor localizing robots should consider incorporating source distance estimation algorithms to improve their search efficiency.

## MATERIALS AND METHODS

### Experimental Setup

#### Black Rock desert setup

Our experimental setup consisted of two components, the first being the mobile sensor stack that was carried by a human for collecting odor signals, and the second component included the placement of 8 stationary wind sensors for ambient wind measurements around the odor source. The odor source was a propylene gas tank (Fig. 1a) that was mounted with a stationary GPS antenna which sent Real Time Kinematics (RTK) correction data to the mobile sensor stack (ublox ANN-MB-00-00 Multiband GNSS Antenna connected to a ublox ZED-F9P receiver attached with an RN41 Bluetooth module for pairing with a Dell Laptop running ROS in Ubuntu Linux OS), which together provided high-resolution position with an accuracy close to 1cm.

The mobile sensor stack in Fig. 1B included a mini photo ionization detector (201A portable, battery powered miniPID photo-ionization detector from Aurora Scientific, operated at its maximum gain settings providing a gas sampling rate of 1250 SCCM), a GPS antenna (with the same ublox receiver Bluetooth module as above) to receive accurate location measurements, and an IMU that provided angular velocity measurements. The sensor stack was balanced on a gimbal for stability and ease of carrying. The odor sensor data was collected using a data acquisition board (MC Measurement Computing-USB1608GX), which was connected along with all the other sensors to a computer that was running ROS as middleware and recorded data in real time. Due to the different sampling rates of sensors (wind sensors: 40 Hz, GPS: 5 Hz), the data were linearly interpolated with respect to the rate of the odor sensor which sampled at 200 Hz. Around the odor source, 7 ambient wind sensors (LI-550F TriSonica Mini Wind and Weather Sensor, Trisonica/Li-Cor) were placed 2 meters above the ground in a square fashion approximately 30 meters away from the source, and one other wind sensor was placed 1 meter away from the odor source for accurate measurement of wind speed and direction near the source as in Fig. 1A, C.

#### Whittell forest setup

A similar arrangement of odor source and wind sensors was set up in Whittell Forest to collect odor encounter data under a forest canopy as in Fig. 2A and 2B. The array of sensors was placed in an L-shaped fashion to better sample the heterogenous environment (Fig. 2C).

### Whiff statistics for trajectory snippets

To understand how a probability distribution of whiff statistics from a short time history is correlated with distance, we randomly selected trajectory snippets of a given length of time. In some cases, our trajectory snippets spanned a wide range of distances. To minimize the artifacts such snippets would create, we first split the datasets from each of our three scenarios into three distance classes (0-5m, 5-30m, and >=30 m). We randomly selected a total of 500 random trajectory snippets from each distance class, ensuring that each trajectory snippet was unique to a single region.

### Calculating whiff frequency (wf)

Whiff frequency is a representation of how often a whiff was encountered within a given duration of time. Following previous applications of whiff frequency in related studies, we used a 1 second time window for calculating whiff frequency [32]. We converted the time series of odor signals to a binary sequence of delta functions, resulting in a time trace that was zero everywhere except for the first frame of each whiff which were assigned a value of 1, see Fig. 4*E*_(*i*)_. We convolved this sequence of whiff onsets with an exponential filter to determine the whiff frequency:

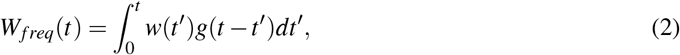

where *w*(*t*^*′*^) is the binary whiff onset signal, *t*^*′*^ is the time at which we are calculating the whiff frequency. The kernel function *g* is defined as follows,

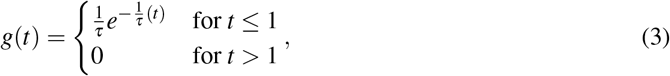

where τ is the decay constant, a measure of how long the previous whiff should be remembered. Based on the whiff data and time constants chosen in [32], we chose τ = 2 sec (Fig. 4 *D*_(*ii*)_). To calculate a single whiff frequency value for each whiff, we averaged the whiff frequency across each individual whiff.

### Kalman Smoothing

We calculated a continuous estimate of distance to the odor source using a Kalman smoother [47]. We defined a linear and time-invariant discrete dynamical systems model,

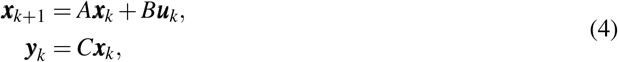

where ***x*** describes two states: the distance to the source, and the velocity towards or away from the source. For simplicity, we did not include any control inputs in our model, instead accounting for this modeling uncertainty in our covariance matrix *Q*, resulting in a simple constant velocity dynamic model,

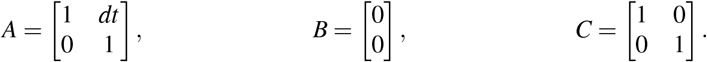

In a real-world application of an agent searching for an odor source, neither of the states we defined can be directly measured (because the source location is unknown). Thus, for our measurements we used the following two (imperfect) observer models to estimate a proxy for these states that are then incorporated into the Kalman smoother. We account for the uncertainty resulting from this discrepancy through our measurement covariance matrix *R*. We calculated an estimate of the distance using our combined regression model (Eqn. 1) to make an intermittent distance prediction associated with each whiff, based on the lookback time of 10-seconds. We also compared the performance of distance estimates when using scenario-specific regression models (Fig. S12). We estimated the velocity towards or away from the source by calculating the sensor’s ground speed component that was parallel to the current ambient wind direction.

For the HWS and LWS scenarios, we defined the measurement and model covariances (*R* and *Q*) to account for the uncertainties in the measurements (*R*) and the dynamic model (*Q*) as follows:

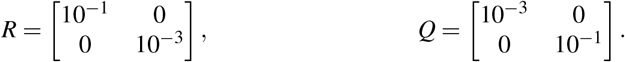

For the Forest scenario, we were able to reduce the median error with slightly different covariance matrices, which help to account for the increase in variability of wind direction in that dataset:

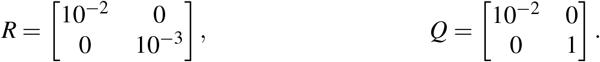

## AUTHOR CONTRIBUTIONS

A.N. and F.v.B. designed research; A.N. and F.v.B. performed research; A.N. and F.v.B. contributed new reagents/analytic tools; A.N. analyzed data; and A.N. and F.v.B. wrote the paper.

## DATA AVAILABILITY

Data is publicly available at: https://doi.org/10.5061/dryad.2547d7wvr.

## CODE AVAILABILITY

All analysis code is available on github (https://github.com/arunavanag591/odor_analysis) and zenodo (https://zenodo.org/doi/10.5281/zenodo.11212243.

## ACKNOWLEDGMENTS

We thank David Stupski and UNR’s StatsChats group for helpful discussions about our statistical analysis, and Jeff Riffell, Kathy Nagel, and Dennis Mathew for feedback on the manuscript. We also thank Jaleesa Houle for her assistance with data collection at the Whittell Forest.

## Supplemental Information

## SUPPLEMENTAL TABLES

**Supplementary Table S1.** Number of whiffs for each environment when the odor signal is thresholded at 4.5 a.u. The original odor signal (sampled at 200Hz) is filtered by a 2nd order butterworth low pass filter to calculate the number of whiffs at 90Hz.

| Number of Whiffs |  |  |
| --- | --- | --- |
| Dataset | 90 Hz | 200 Hz |
| HWS | 2760 | 2842 |
| LWS | 3517 | 3522 |
| Forest | 4830 | 4986 |

**Supplementary Table S2.** Comparison of ***R***^**2**^ and relative motion adjusted ***R***^**2**^ for individual whiff statistics when correlated with distance from source. To account for relative motion, we first determined the relationship between each whiff statistic and the sensor’s velocity parallel and perpendicular to the wind: *w****c*** ∼ *v*_||_ + *v*_∼_. In our regression with source distance, we then used the residuals for each whiff statistic after removing the relative motion effects: *w****c***_***r***_ **=** *w****c* ∼ *α*** _**1**_ **∼ *v***_||_ **∼ *α*** _**2**_ **∼ *v***_∼_.

| Dataset | Original $R^2$ | Relative Motion Adjusted $R^2$ |
| --- | --- | --- |
| HWS | 0.14 | 0.14 |
| LWS | 0.014 | 0.13 |
| Forest | 0.08 | 0.07 |
| All Combined | 0.13 | 0.13 |

**Supplementary Table S3.**
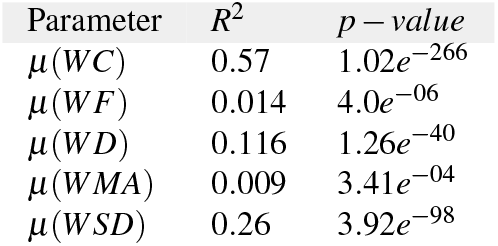
***R***^**2**^ and p-value for individual meta whiff statistics for a lookback window of 10 seconds, correlated with distance from source.

| Parameter | $R^2$ | p-value |
| --- | --- | --- |
| $\mu(WC)$ | 0.57 | $1.02e^{-266}$ |
| $\mu(WF)$ | 0.014 | $4.0e^{-06}$ |
| $\mu(WD)$ | 0.116 | $1.26e^{-40}$ |
| $\mu(WMA)$ | 0.009 | $3.41e^{-04}$ |
| $\mu(WSD)$ | 0.26 | $3.92e^{-98}$ |

**Supplementary Table S4.** ***AIC*** and **Δ*AIC*** for selecting the most significant statistics that correlate with distance from source.

| $min(AIC)$ | $\Delta AIC$ | Parameters |
| --- | --- | --- |
| 10726.2 | 0 | 0 |
| 10655.9 | -70.3 | $\mu(WC)$ |
| 10631.2 | -24.7 | $\mu(WC), \sigma(WMA)$ |
| 10594.5 | -36.6 | $\mu(WC), \sigma(WMA), max(WMA)$ |
| 10575.1 | -19.5 | $\mu(WC), \sigma(WMA), max(WMA), \sigma(WD)$ |
| 10557.3 | -17.8 | $\mu(WC), \sigma(WMA), max(WMA), \sigma(WD), max(WC)$ |

**Supplementary Table S5.** ***AIC*** and **Δ*AIC*** for selecting the most significant statistics that correlate with distance from source (10 iterations).

| $min(AIC)$ | $\Delta AIC$ | Parameters |
| --- | --- | --- |
| 10726.2144 | 0 | 0 |
| 10655.8883 | -70.3260 | $\mu(WC)$ |
| 10631.1534 | -24.7349 | $\mu(WC), \sigma(WMA)$ |
| 10594.5044 | -36.6490 | $\mu(WC), \sigma(WMA), max(WMA)$ |
| 10575.0510 | -19.4534 | $\mu(WC), \sigma(WMA), max(WMA), \sigma(WD)$ |
| 10557.2739 | -17.7770 | $\mu(WC), \sigma(WMA), max(WMA), \sigma(WD), max(WC)$ |
| 10532.4997 | -24.7742 | $\mu(WC), \sigma(WMA), max(WMA), \sigma(WD), max(WC), min(WC)$ |
| 10529.4179 | -3.0818 | $\mu(WC), \sigma(WMA), max(WMA), \sigma(WD), max(WC), min(WC), WC_k$ |
| 10524.6016 | -4.8162 | $\mu(WC), \sigma(WMA), max(WMA), \sigma(WD), max(WC), min(WC), WC_k, \mu(WMA),$ |

## SUPPLEMENTAL FIGURES

**Supplementary Figure S1.**
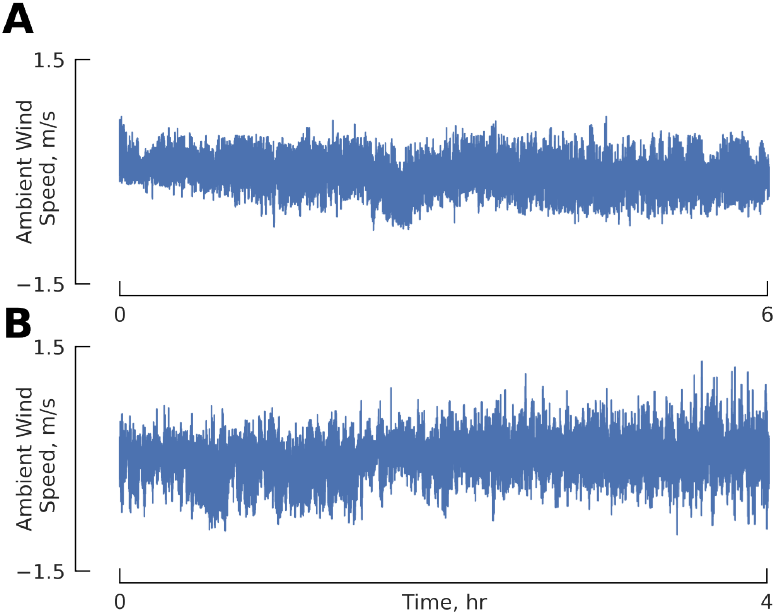
Vertical ambient wind velocity remained small and mostly constant throughout the experiment period in **A**. the Desert and **B**. the Forest.

**Supplementary Figure S2.**
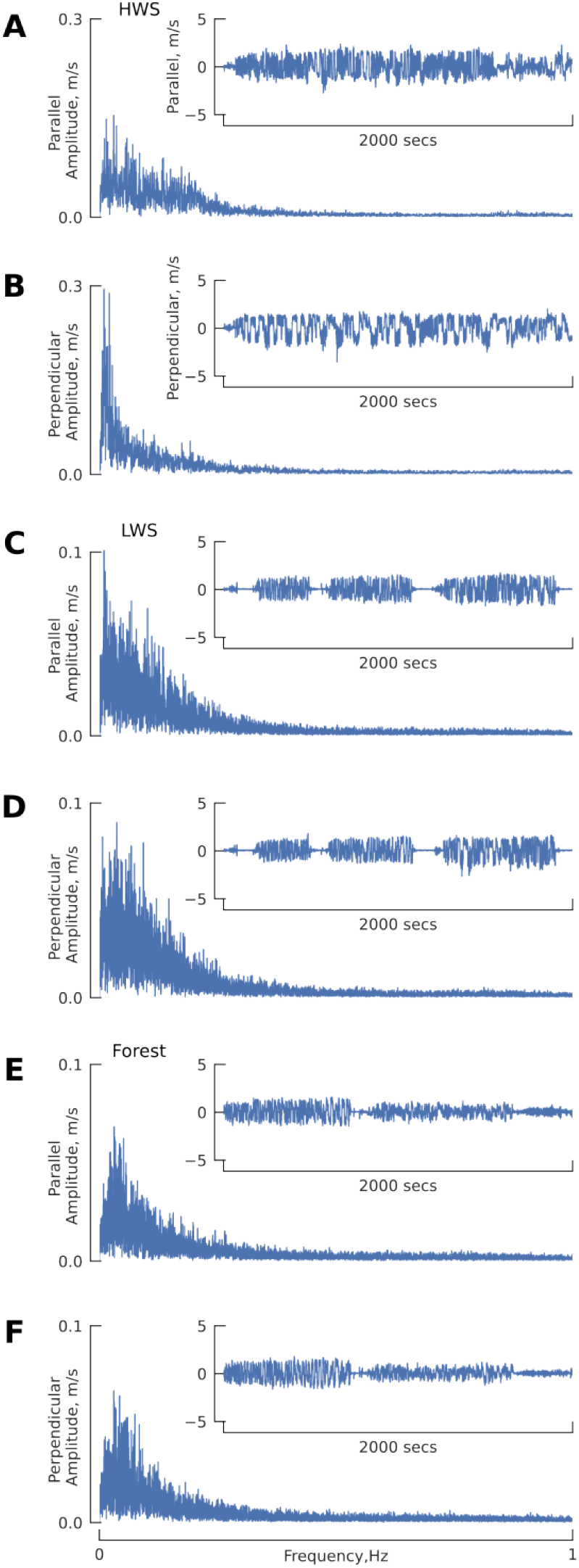
Sensor ground velocity (from GPS measurements) projected onto the unit vector pointed towards the source (parallel component) or perpendicular to that direction. Insets show raw time series plots of these velocities, and the larger plots show the FFT of each component. **A-B** HWS, **C-D** LWS, **E-F** Forest datasets. Dynamics similar to casting are best seen in the HWS scenario, where we made more frequent movements perpendicular to the direction pointing towards the source. In the LWS and forest scenarios we made more randomized movements to find the odor plume.

**Supplementary Figure S3.**
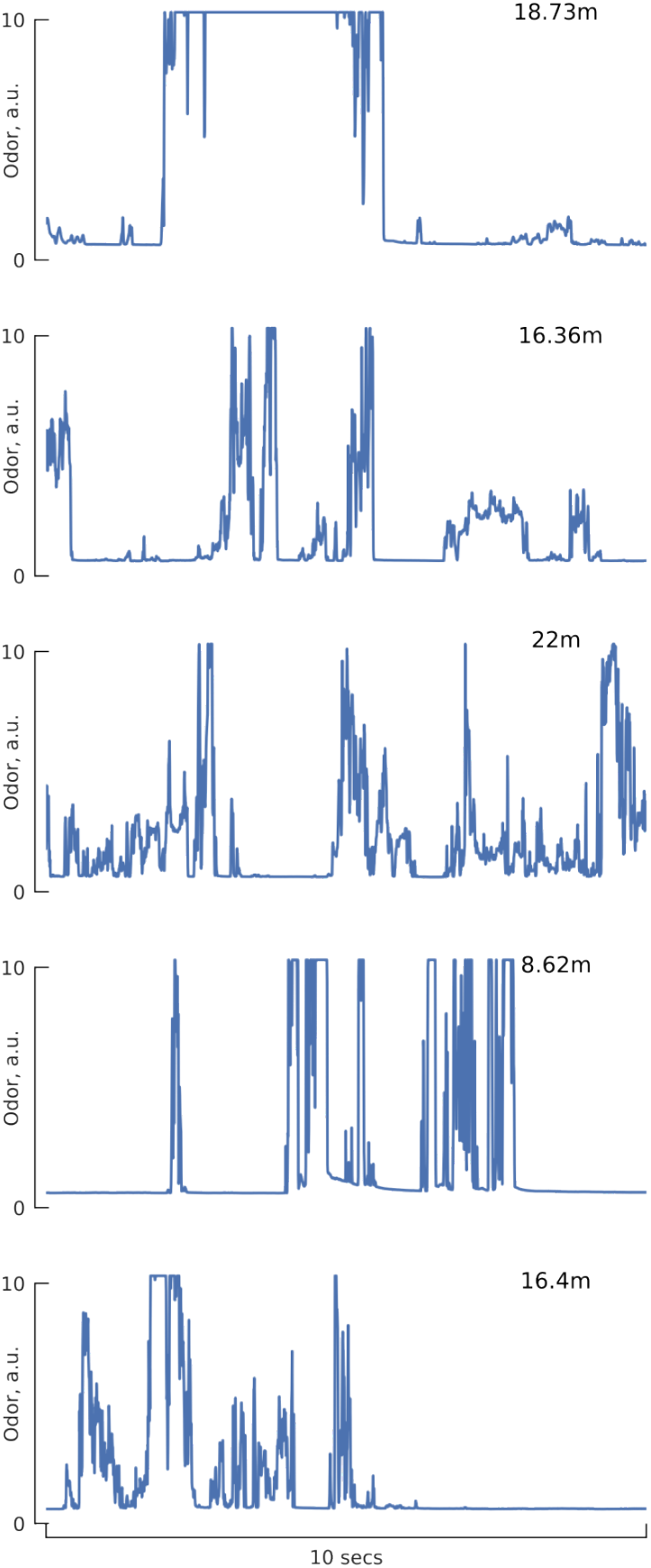
Examples of 10 second odor signal traces, illustrating the intermittent nature of odor in the wind. Numbers in upper right corners show the average distance of the encounters from the source.

**Supplementary Figure S4.**
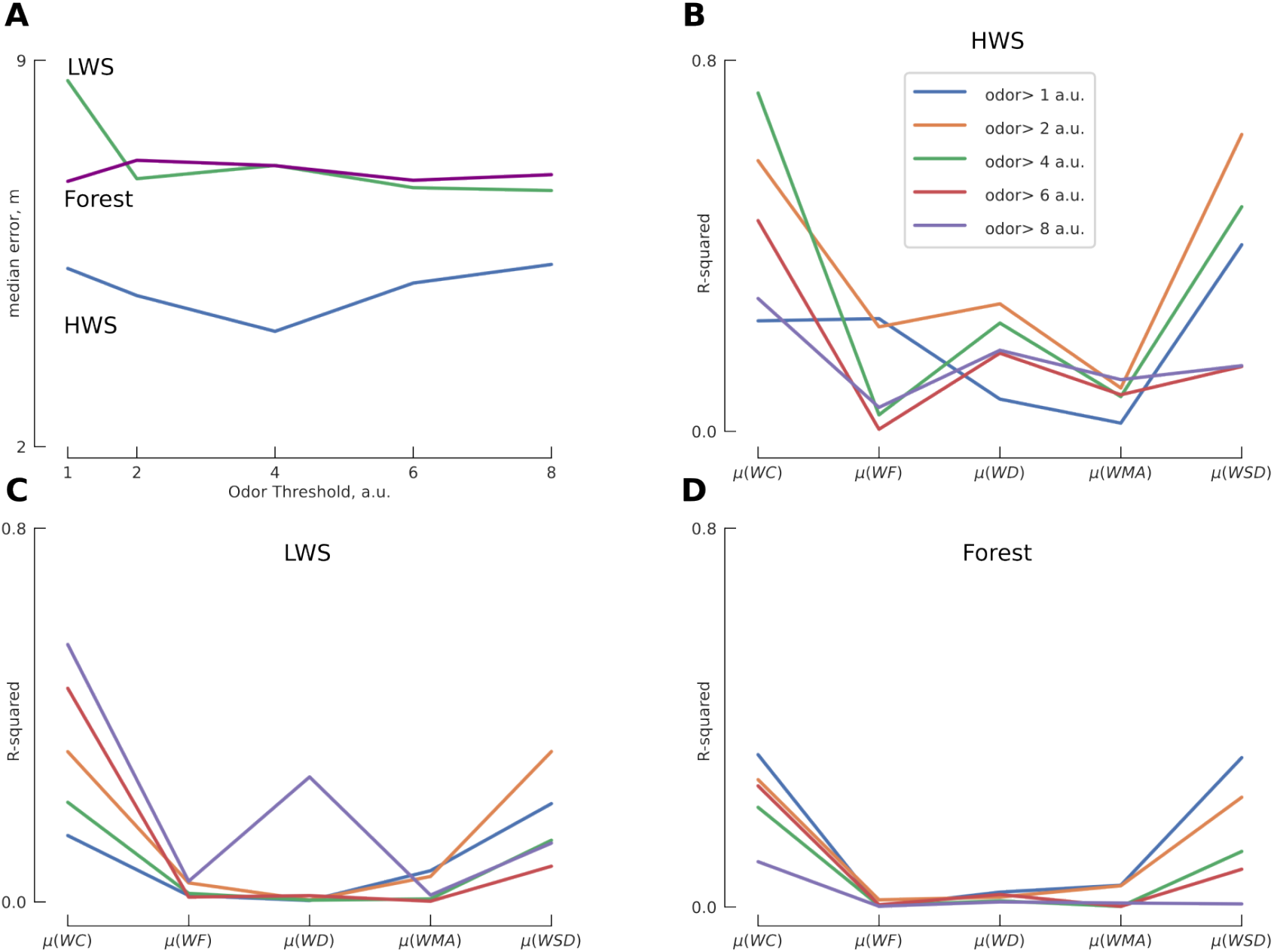
Sensitivity analysis of key conclusions with respect to choice of odor threshold. For the analysis presented in the main text we chose a odor threshold of 4.5 a.u. **A**. Median error of continuous estimates of source distance using our Kalman smoother approach (following Fig. 7) for different choices of odor thresholds. **B-D**. Shows the *R*^2^ value for mean whiff statistics for a lookback window of 10 seconds correlated with source distance (following Table S3. across a range of odor thresholds for HWS, LWS, and Forest.

**Supplementary Figure S5.**
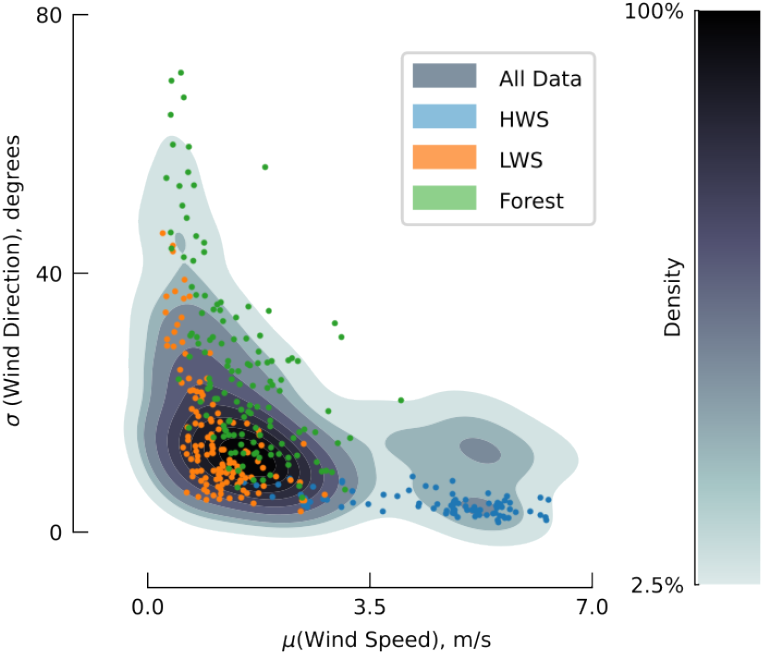
Wind characteristics from our LWS, HWS and Forest scenarios span a representative range of near surface wind characteristics from a larger dataset. We characterized the wind by calculating the standard deviation of direction and mean wind speed for equally spaced 10 second windows. The contour shows the distribution of these wind characteristics from 10 individual days of wind measurements in playa, sage steppe, and forest environments from [1]. The scatter plot shows these wind characteristics for the LWS, HWS and Forest scenarios. The larger dataset includes the data from the LWS, HWS, and Forest scenarios in addition to 8 other days of data collection.

**Supplementary Figure S6.**
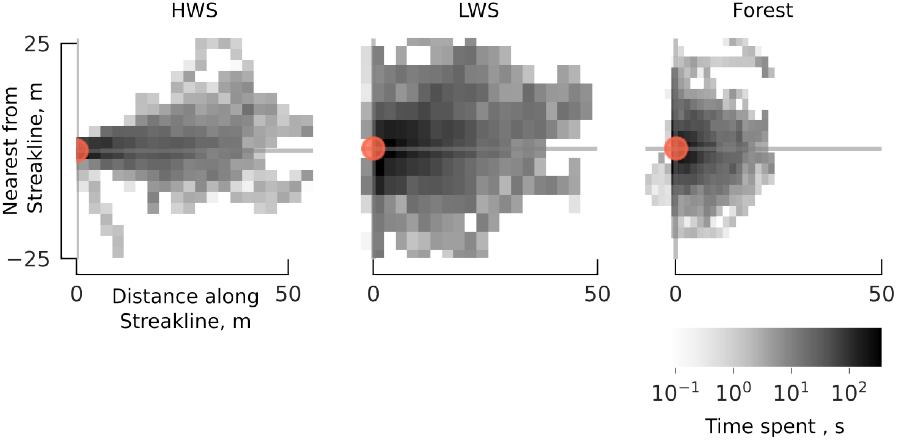
Using the coordinate system introduced in Fig. 3B, this figure shows a representation of how much time the sensor was in each location relative to the streakline and odor source.

**Supplementary Figure S7.**
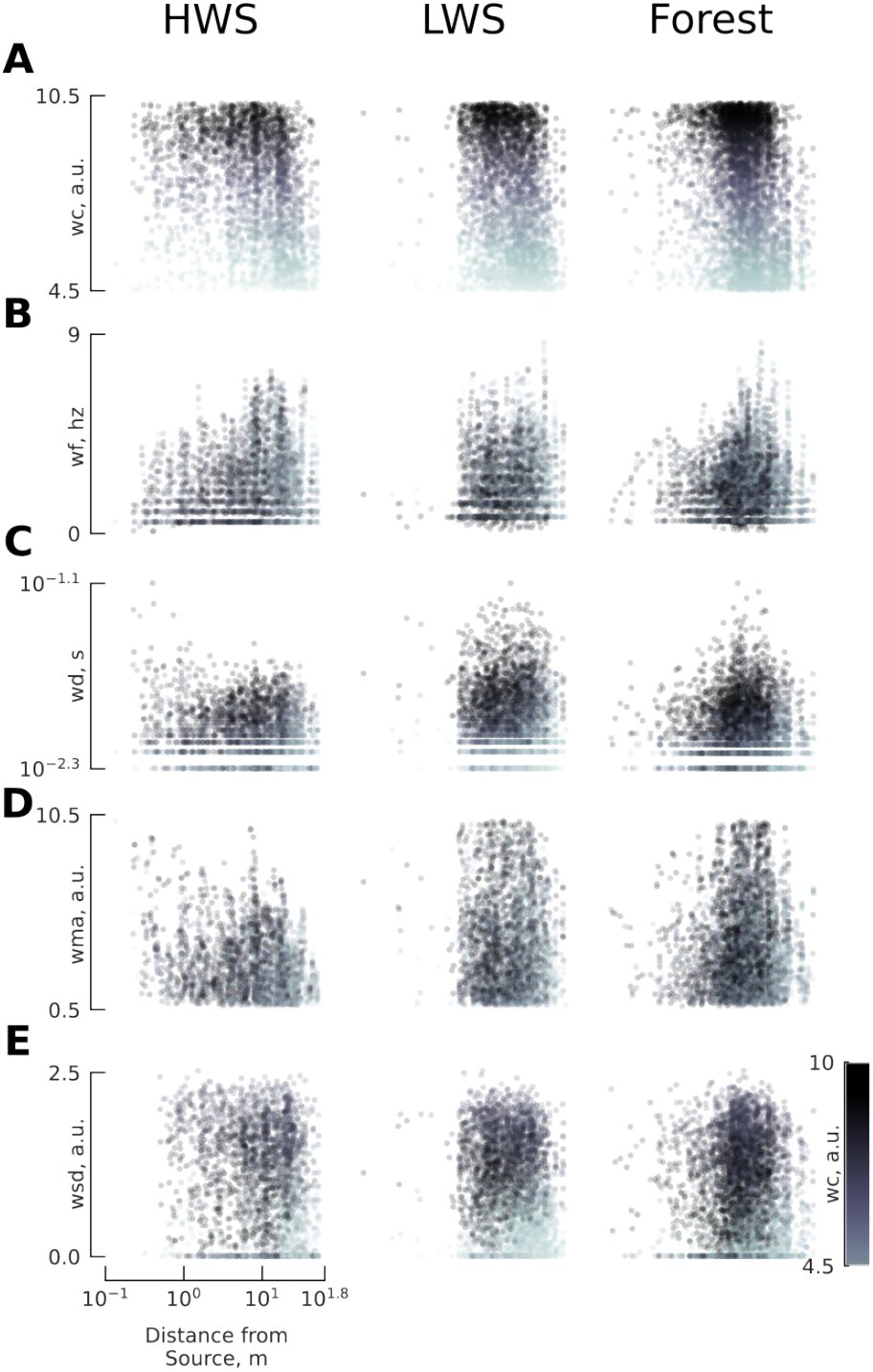
Individual whiff statistics vs distance from source for HWS, LWS and Forest (column wise) **A**. Whiff concentration **B**. Whiff frequency, **C**. Whiff duration, **D**. Whiff moving average **E**. Whiff standard deviation.

**Supplementary Figure S8.**
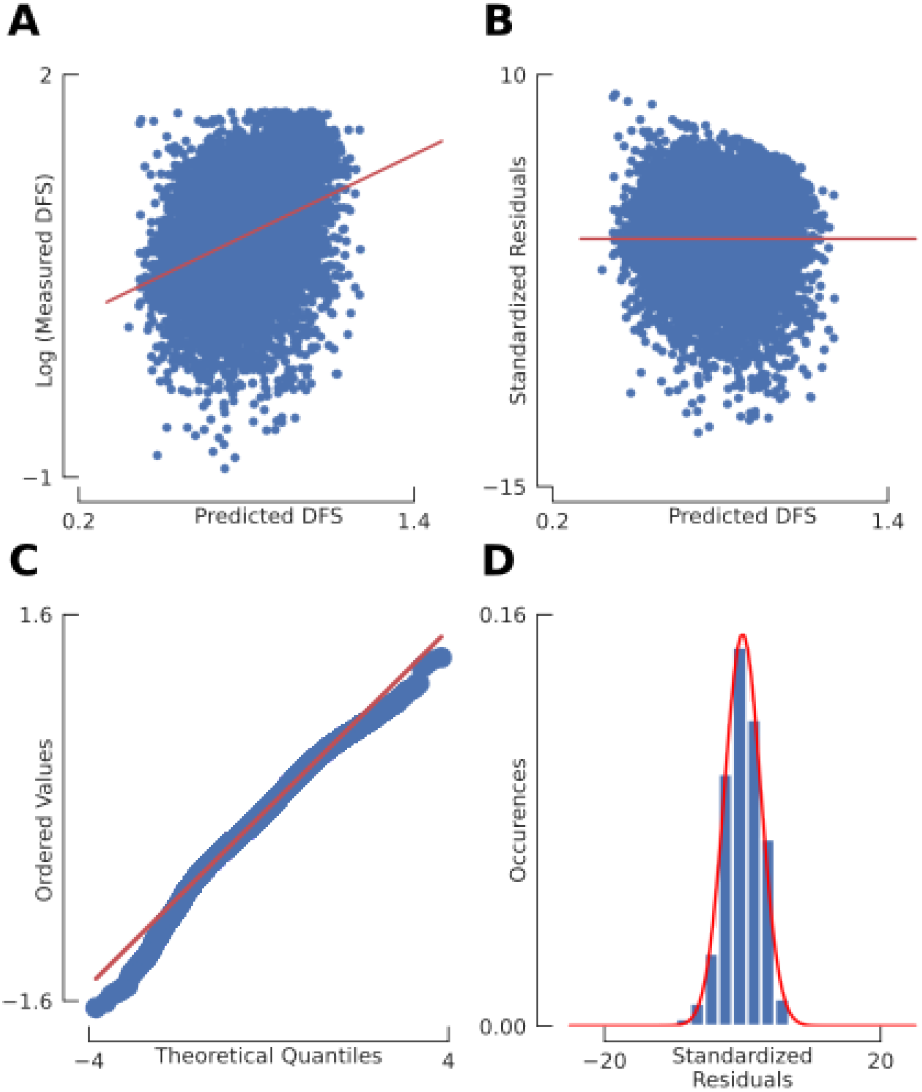
Comprehensive residual analysis for model validation **A**. Actual vs predicted. Scatter plot illustrating the correlation between the actual and predicted values, providing an overview of the model’s accuracy. **B**. Standardized residuals. A plot of the standardized residuals helps identify any outliers that could potentially influence the model’s performance. **C**. Residual analysis. Q-Q plot: A quantile-quantile plot used to assess if the residuals follow a normal distribution. **D**. Normality of residuals. A histogram with an overlaid normal curve to visually inspect the distribution of the residuals.

**Supplementary Figure S9.**
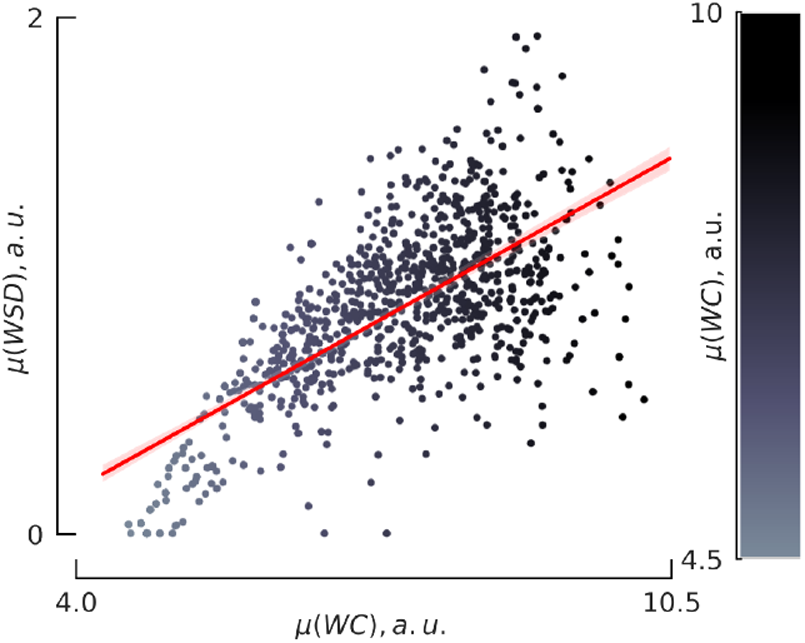
Relationship between meta statistics for the mean whiff concentration (*µ*(*WC*)) and mean whiff standard deviation (*µ*(*WSD*)). The trend line shows they are almost linearly correlated. This helps to explain why the △*AIC* analysis in Table S4 did not select *µ*(*WSD*), even though it has the second highest correlation with distance from the source (Fig. S10).

**Supplementary Figure S10.**
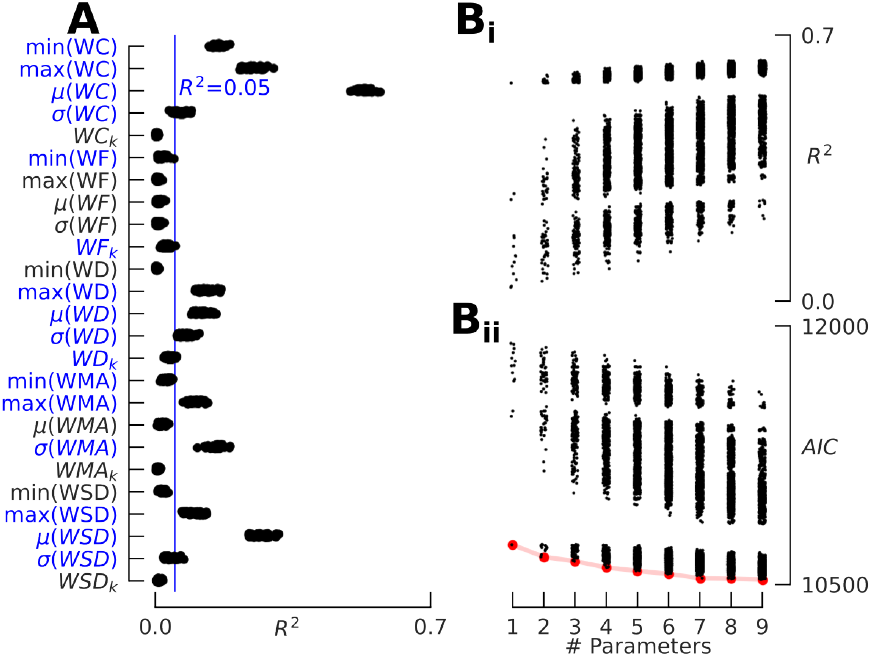
A. Bootstrapped *R*^2^ is shown for each meta statistic. For subsequent analyses, we focus on meta-statistics that reached above a threshold of *R*^2^ = 0.05 (blue). **B**. To find the most important meta statistics, we calculated the *R*^2^ and *AIC* for all possible models containing up to 9 meta statistics. The red line in **B**_**ii**_ shows the best models.

**Supplementary Figure S11.**
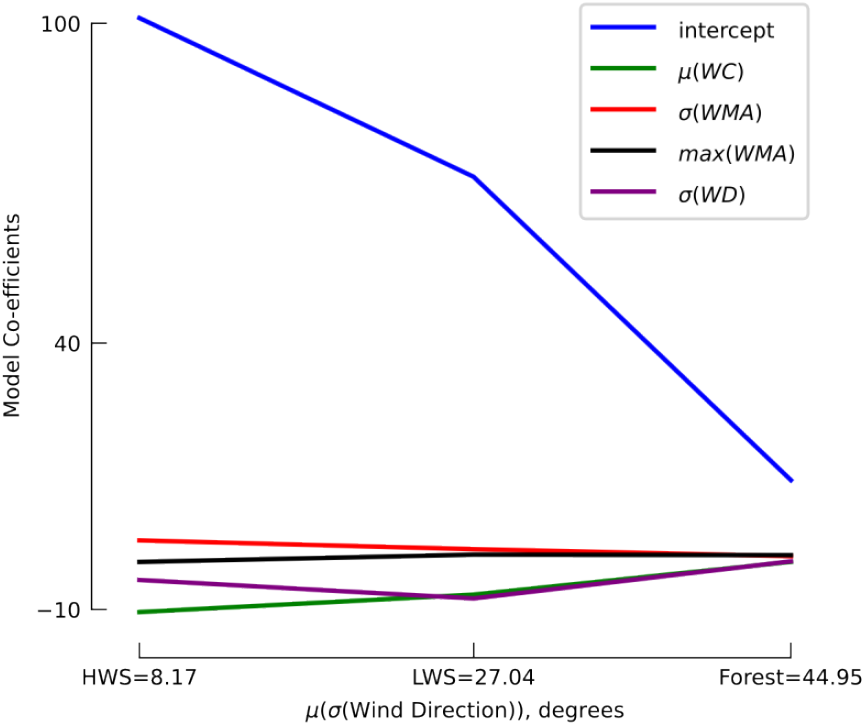
The *AIC* filtered parameter coefficients are similar across all three of our scenarios, except for the intercept values. The x-axis shows the mean of the standard deviations of the wind direction for each environment that are shown in Fig. S5.

**Supplementary Figure S12.**
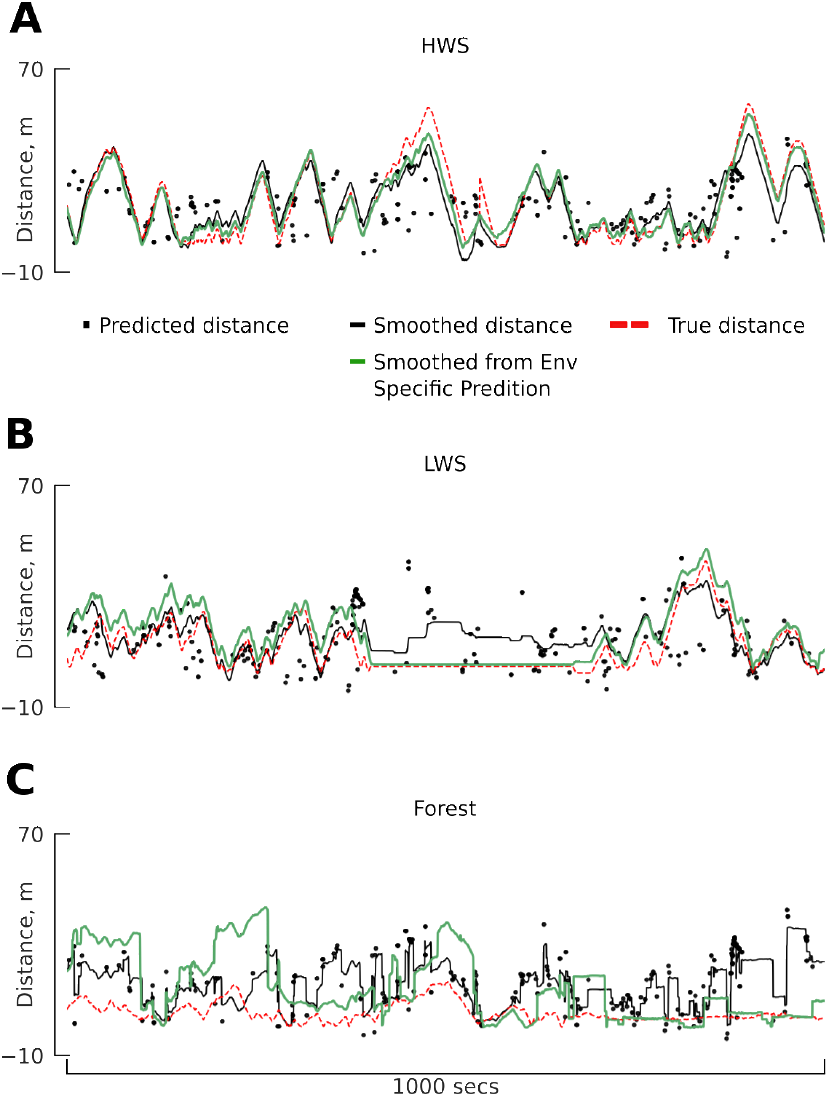
Kalman smoothed estimates of the distance to the odor source (red and black lines) vs intermittent distance predictions from our 4-parameter regression model for the 10-second lookback window (black dots) vs true distance (red dashed line), similar to Fig. 7. The black lines (copied from Fig. 7) shows the smoothed distance estimates when using the same combined model for all three scenarios (distance∼ model). The green lines show the smoothed distance estimates when environment-specific regression models are used: **A** HWS (distance = -10.74 *µ*(WC) - 0.63 max (WMA) + 1.16 *σ* (WMA) - 7.45 *σ* (WD) + 102.47) **B**. LWS (distance = -7.59 *µ*(WC) + 0.52 max(WMA) - 0.82 *σ* (WMA) - 7.92 *σ* (WD) + 73.24) **C**. Forest (distance = -1.3 *µ*(WC) + 0.09 max(WMA) - 0.06 *σ* (WMA) - 0.53 *σ* (WD) + 15.05).

## REFERENCES

[1] Baker, K. L. et al. Algorithms for olfactory search across species. Journal of Neuroscience 38, 9383–9389 (2018).

[2] Chen, X.-x. & Huang, J. Odor source localization algorithms on mobile robots: A review and future outlook. Robotics and Autonomous Systems 112, 123–136 (2019).

[3] Jing, T., Meng, Q.-H. & Ishida, H. Recent progress and trend of robot odor source localization. IEEJ Transactions on Electrical and Electronic Engineering 16, 938–953 (2021).

[4] Murlis, J., Willis, M. A. & Cardé, R. T. Spatial and temporal structures of pheromone plumes in fields and forests. Physiological entomology 25, 211–222 (2000).

[5] Riffell, J. A., Abrell, L. & Hildebrand, J. G. Physical processes and real-time chemical measurement of the insect olfactory environment. Journal of chemical ecology 34, 837–853 (2008).

[6] Connor, E. G., McHugh, M. K. & Crimaldi, J. P. Quantification of airborne odor plumes using planar laser-induced fluorescence. Experiments in Fluids 59, 1–11 (2018).

[7] Celani, A., Villermaux, E. & Vergassola, M. Odor landscapes in turbulent environments. Physical Review X 4, 041015 (2014).

[8] van Breugel, F. & Dickinson, M. H. Plume-tracking behavior of flying drosophila emerges from a set of distinct sensory-motor reflexes. Current Biology 24, 274–286 (2014).

[9] Kanzaki, R., Sugi, N. & Shibuya, T. Self-generated zigzag turning of bombyx mori males during pheromone-mediated upwind walking. Zoological science 9, 515–527 (1992).

[10] Vickers, N. J. & Baker, T. C. Reiterative responses to single strands of odor promote sustained upwind flight and odor source location by moths. Proceedings of the National Academy of Sciences 91, 5756–5760 (1994).

[11] Page, J. L., Dickman, B. D., Webster, D. R. & Weissburg, M. J. Getting ahead: context-dependent responses to odorant filaments drive along-stream progress during odor tracking in blue crabs. Journal of Experimental Biology 214, 1498–1512 (2011).

[12] Scholz, A. T., Horrall, R. M., Cooper, J. C. & Hasler, A. D. Imprinting to chemical cues: The basis for home stream selection in salmon. Science 192, 1247–1249 (1976).

[13] Findley, T. M. et al. Sniff-synchronized, gradient-guided olfactory search by freely moving mice. Elife 10, e58523 (2021).

[14] Farrell, J. A., Pang, S. & Li, W. Chemical plume tracing via an autonomous underwater vehicle. IEEE Journal of Oceanic Engineering 30, 428–442 (2005).

[15] Hutchinson, M., Liu, C. & Chen, W.-H. Information-based search for an atmospheric release using a mobile robot: Algorithm and experiments. IEEE Transactions on Control Systems Technology 27, 2388–2402 (2018).

[16] Shigaki, S., Yoshimura, Y., Kurabayashi, D. & Hosoda, K. Palm-sized quadcopter for three-dimensional chemical plume tracking. IEEE Transactions on Instrumentation and Measurement 71, 1–12 (2022).

[17] Anderson, M. J., Sullivan, J. G., Horiuchi, T. K., Fuller, S. B. & Daniel, T. L. A bio-hybrid odor-guided autonomous palm-sized air vehicle. Bioinspiration & Biomimetics 16, 026002 (2020).

[18] Anderson, M. J. et al. The “smellicopter,” a bio-hybrid odor localizing nano air vehicle. In 2019 IEEE/RSJ International Conference on Intelligent Robots and Systems (IROS), 6077–6082 (IEEE, 2019).

[19] Luo, B., Meng, Q.-H., Wang, J.-Y. & Zeng, M. A flying odor compass to autonomously locate the gas source. IEEE Transactions on Instrumentation and Measurement 67, 137–149 (2017).

[20] Shigaki, S., Yamada, M., Kurabayashi, D. & Hosoda, K. Robust moth-inspired algorithm for odor source localization using multimodal information. Sensors 23, 1475 (2023).

[21] Pang, R., van Breugel, F., Dickinson, M., Riffell, J. A. & Fairhall, A. History dependence in insect flight decisions during odor tracking. PLoS computational biology 14, e1005969 (2018).

[22] Grünbaum, D. & Willis, M. A. Spatial memory-based behaviors for locating sources of odor plumes. Movement ecology 3, 1–21 (2015).

[23] Saxena, N., Natesan, D. & Sane, S. P. Odor source localization in complex visual environments by fruit flies. Journal of Experimental Biology 221, jeb172023 (2018).

[24] Raguso, R. A. & Willis, M. A. Synergy between visual and olfactory cues in nectar feeding by naive hawkmoths, manduca sexta. Animal Behaviour 64, 685–695 (2002).

[25] Jinn, J., Connor, E. G. & Jacobs, L. F. How ambient environment influences olfactory orientation in search and rescue dogs. Chemical Senses 45, 625–634 (2020).

[26] Wechsler, S. P. & Bhandawat, V. Behavioral algorithms and neural mechanisms underlying odor-modulated locomotion in insects. Journal of Experimental Biology 226 (2023).

[27] Rigolli, N., Magnoli, N., Rosasco, L. & Seminara, A. Learning to predict target location with turbulent odor plumes. eLife 11, e72196 (2022).

[28] Yee, E., Kosteniuk, P. R., Chandler, G. M., Biltoft, C. A. & Bowers, J. F. Statistical characteristics of concentration fluctuations in dispersing plumes in the atmospheric surface layer. Boundary-Layer Meteorology 65, 69–109 (1993).

[29] Boie, S. D. et al. Information-theoretic analysis of realistic odor plumes: What cues are useful for determining location? PLoS computational biology 14, e1006275 (2018).

[30] Park, I. J. et al. Neurally encoding time for olfactory navigation. PLOS Computational Biology 12, e1004682 (2016).

[31] Schmuker, M., Bahr, V. & Huerta, R. Exploiting plume structure to decode gas source distance using metal-oxide gas sensors. Sensors and Actuators B: Chemical 235, 636–646 (2016).

[32] Demir, M., Kadakia, N., Anderson, H. D., Clark, D. A. & Emonet, T. Walking drosophila navigate complex plumes using stochastic decisions biased by the timing of odor encounters. Elife 9, e57524 (2020).

[33] Kadakia, N. et al. Odour motion sensing enhances navigation of complex plumes. Nature 1–8 (2022).

[34] Jayaram, V., Kadakia, N. & Emonet, T. Sensing complementary temporal features of odor signals enhances navigation of diverse turbulent plumes. eLife 11, e72415 (2022).

[35] Zocchi, D., Emily, S. Y., Hauser, V., O’Connell, T. F. & Hong, E. J. Parallel encoding of co2 in attractive and aversive glomeruli by selective lateral signaling between olfactory afferents. Current Biology 32, 4225–4239 (2022).

[36] álvarez-Salvado, E. et al. Elementary sensory-motor transformations underlying olfactory navigation in walking fruit-flies. Elife 7, e37815 (2018).

[37] Houle, J. & van Breugel, F. Near-surface wind variability over spatiotemporal scales relevant to plume tracking insects. Physics of Fluids 35 (2023).

[38] van Breugel, F., Jewell, R. & Houle, J. Active anemosensing hypothesis: how flying insects could estimate ambient wind direction through sensory integration and active movement. Journal of the Royal Society Interface 19, 20220258 (2022).

[39] Willis, M. & Arbas, E. Odor-modulated upwind flight of the sphinx moth, manduca sexta l. Journal of Comparative Physiology A 169 (1991).

[40] Akaike, H. A new look at the statistical model identification. IEEE transactions on automatic control 19, 716–723 (1974).

[41] Butterworth, S. On the theory of filter amplifiers. Wireless Engineer 7, 536–541 (1930).

[42] Van Breugel, F., Riffell, J., Fairhall, A. & Dickinson, M. H. Mosquitoes use vision to associate odor plumes with thermal targets. Current Biology 25, 2123–2129 (2015).

[43] Bidlingmayer, W. & Hem, D. The range of visual attraction and the effect of competitive visual attractants upon mosquito (diptera: Culicidae) flight. Bulletin of Entomological Research 70, 321–342 (1980).

[44] Nagel, K. I. & Wilson, R. I. Biophysical mechanisms underlying olfactory receptor neuron dynamics. Nature Neuroscience 14, 208–216 (2011).

[45] Egea-Weiss, A., Renner, A., Kleineidam, C. J. & Szyszka, P. High precision of spike timing across olfactory receptor neurons allows rapid odor coding in drosophila. iScience 4, 76–83 (2018).

[46] Szyszka, P., Gerkin, R. C., Galizia, C. G. & Smith, B. H. High-speed odor transduction and pulse tracking by insect olfactory receptor neurons. Proceedings of the National Academy of Sciences 111, 16925–16930 (2014).

[47] Crassidis, J. L. & Junkins, J. L. Optimal estimation of dynamic systems (CRC press, 2011).

## REFERENCES

[1] Houle, J. & van Breugel, F. Near-surface wind variability over spatiotemporal scales relevant to plume tracking insects. Physics of Fluids 35 (2023).

